# Parallel construction of object motion from retina to cortex

**DOI:** 10.64898/2026.07.27.740872

**Authors:** Javier C. Weddington, David D. Au, Joshua B Melander, Youssef Faragalla, Zaki Alaoui, Stephen A. Baccus

**Author notes:** Correspondence: *Stephen A. Baccus. convergent-oms.github.io.

## Abstract

Detecting a moving object against a moving background is fundamental to survival. The retina is known to perform this computation, extracting object motion from global motion before signals reach the brain. Under the prevailing hierarchical view of vision, downstream areas should inherit and elaborate this retinal computation rather than rebuild it. Using multielectrode recordings of matched stimuli in mouse retina and cortex along with computational modeling, we show instead that post-retinal processing recomputes rather than inherits object motion sensitivity. Retinal and cortical object-motion sensitivities have distinct tuning: the retina prefers fine jitter, whereas cortex prefers coarse drift, revealing a division of labor in which the retina detects moving objects, whereas the cortex conveys information about object pattern. Vision synthesizes object motion through parallel specialization, not hierarchical refinement.

## Introduction

For an animal that must catch prey or avoid becoming it, few visual tasks matter more than spotting and identifying an object moving against a patterned background. Motion is often the only cue available, since a stationary object may be difficult to distinguish from its surroundings. Yet motion is itself ambiguous: because the eyes and head are almost never still, the whole scene sweeps across the retina even during fixation, so an object’s local motion must be separated from this global, self-generated motion (1).

The first stage of the solution is known to lie in the retina. A class of retinal ganglion cells fires vigorously when the receptive-field center moves differently from its surround, yet is suppressed when center and surround move together, as they do during self-motion (1). This object-motion-sensitive (OMS) response is conserved across species (2, 3), is largely invariant to the pattern being moved, and arises from a well-characterized circuit in which the ganglion cell pools over rectified bipolar terminals under inhibition from wide-field amacrine cells (4–8). The retina thus flags moving objects before the signal leaves the eye.

How this signal is then transmitted to higher levels is less clear. Under the prevailing hierarchical view of vision, cortical areas inherit the computations performed upstream and progressively elaborate them rather than rebuilding them (9– 11). In this framework, cortical object-motion sensitivity should be a refined descendant of the retinal OMS signal. Yet primary visual cortex also possesses machinery — orientation selectivity (12), surround suppression (13), and motion integration (14) — that could in principle compute object motion on its own. Here we compare object motion sensitivity in the mouse at the level of the retina and primary visual cortex, using in vivo eye tracking to reconstruct the retinal image so that both stages are probed with matched stimuli. A challenge in the study of visual phenomena is to explore a sufficient range of stimuli in a finite experiment. To address this challenge, we fit convolutional neural network models to each stage (15–24) using a set of natural movies, and used these models to probe a wide range of stimulus conditions beyond those tested experimentally, generating specific predictions about retinal and cortical object-motion sensitivity that we then confirmed with targeted experiments. We find that the two stages are tuned to different scales of motion: the retina to fine jitter, the cortex to coarse drift. Further analysis indicates that the visual cortex does not inherit the retinal computation but recomputes object motion independently. This division of labor — the retina signaling that an object is moving, the cortex conveying pattern information about the object — indicates that the visual system synthesizes object motion through parallel specialization rather than hierarchical refinement.

## Results

We presented a rich, natural visual stimuli to head-fixed mice in vivo and the retina in vitro. Mice ran on a wheel while viewing open-loop, hour-long videos of natural scenes captured at rodent eye level with an omnidirectional camera, spanning optic flow, head movements, translational and rotational motion, abrupt stops, and stationary intervals. We recorded eye position and primary visual cortex (V1) activity with Neuropixel silicon probes in vivo (Fig. 1A, S1). To record retinal responses to the same eye-movement– transformed image seen in vivo, we then reconstructed the gaze-shifted stimulus each animal had actually experienced and replayed it to excised retina on a multielectrode array (Fig. 1C).

**Fig. 1.**
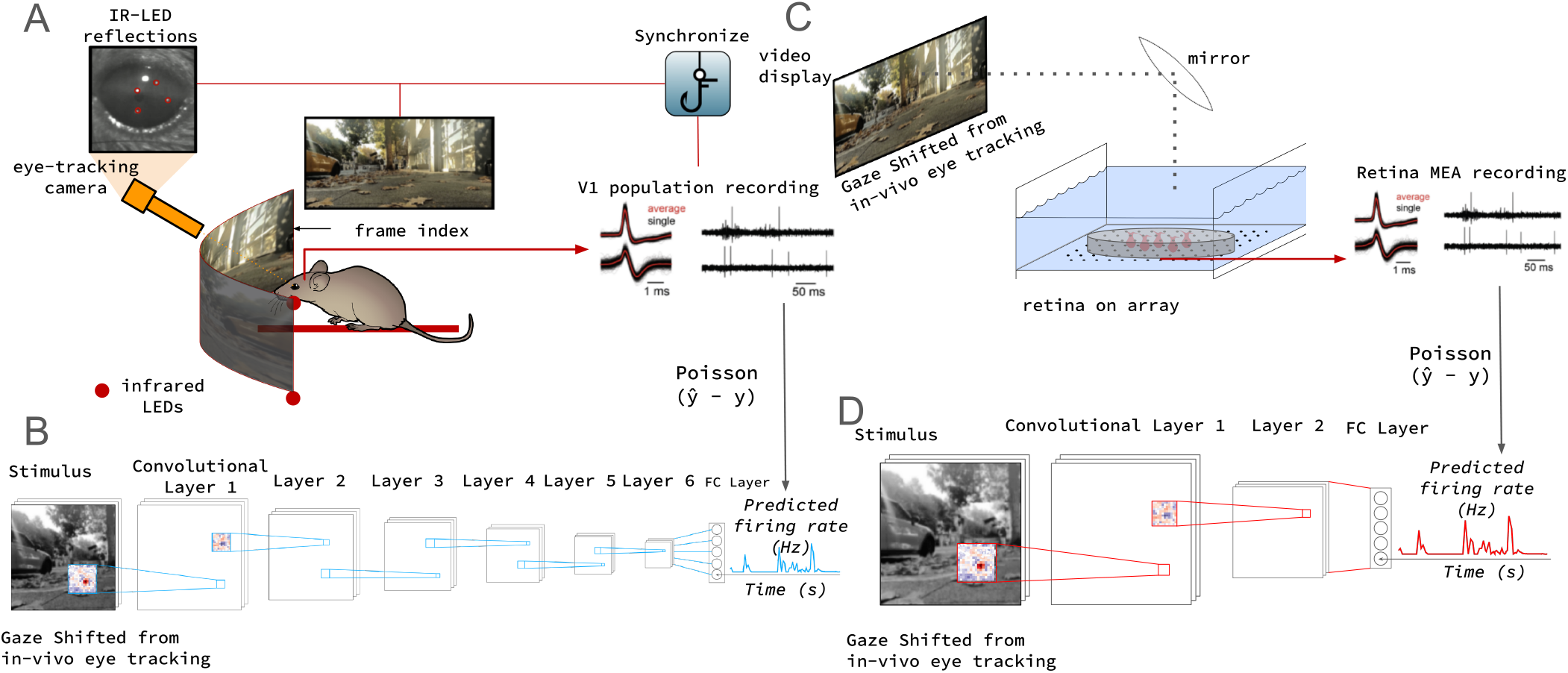
Experimental and modeling pipeline for predicting neural responses to gaze-shifted naturalistic stimuli. **A:** In-vivo V1 recording schematic: A head-fixed mouse views natural scene videos while eye position and V1 Neuropixels activity are simultaneously recorded. **B:** A 6-layer convolutional neural network (CNN) predicts V1 firing rates from gaze-shifted frames using Poisson negative log-likelihood loss (NLLoss). **C:** Ex-vivo setup: The same gaze-shifted stimulus is projected onto an excised retina on a multi-electrode array (MEA). **D:** A 2-layer CNN predicts retinal firing rates from the same stimuli. Model depths reflect cortical versus retinal computational complexity. **Note:** Using identical in-vivo gaze shifts provides matched, behaviorally relevant inputs to directly compare retinal and cortical representations.

We fit separate convolutional neural network (CNN) encoding models to each circuit—a six-layer network for V1 and a three-layer network for retina, depths chosen to reflect the relative computational complexity of the two stages (Fig. 1B,D)—each trained to predict single-unit firing rates from the gaze-shifted stimulus. Models were fit to V1 from 4 mice and to retina from 2 retinas and evaluated on held-out natural-scene trials.

Model fidelity was assessed with three complementary measures (Fig. 2A, Fig. S2A). Split-half reliability estimated the stimulus-driven fraction of each neuron’s variance, setting a model-independent ceiling. Because eye movements differed on every trial, the effective stimulus varied trial to trial; trial-averaged correlation therefore smooths across distinct inputs and provides a conservative lower bound, whereas single-trial correlation, evaluated against each trial’s own gaze-shifted stimulus, measures trial-to-trial predictive fidelity. Beyond these summary statistics, we compared each model unit’s receptive field to the corresponding neuron’s spike-triggered average and found agreement for those cells that had a clear receptive field (Fig. 2C, Fig. S2C).

**Fig. 2.**
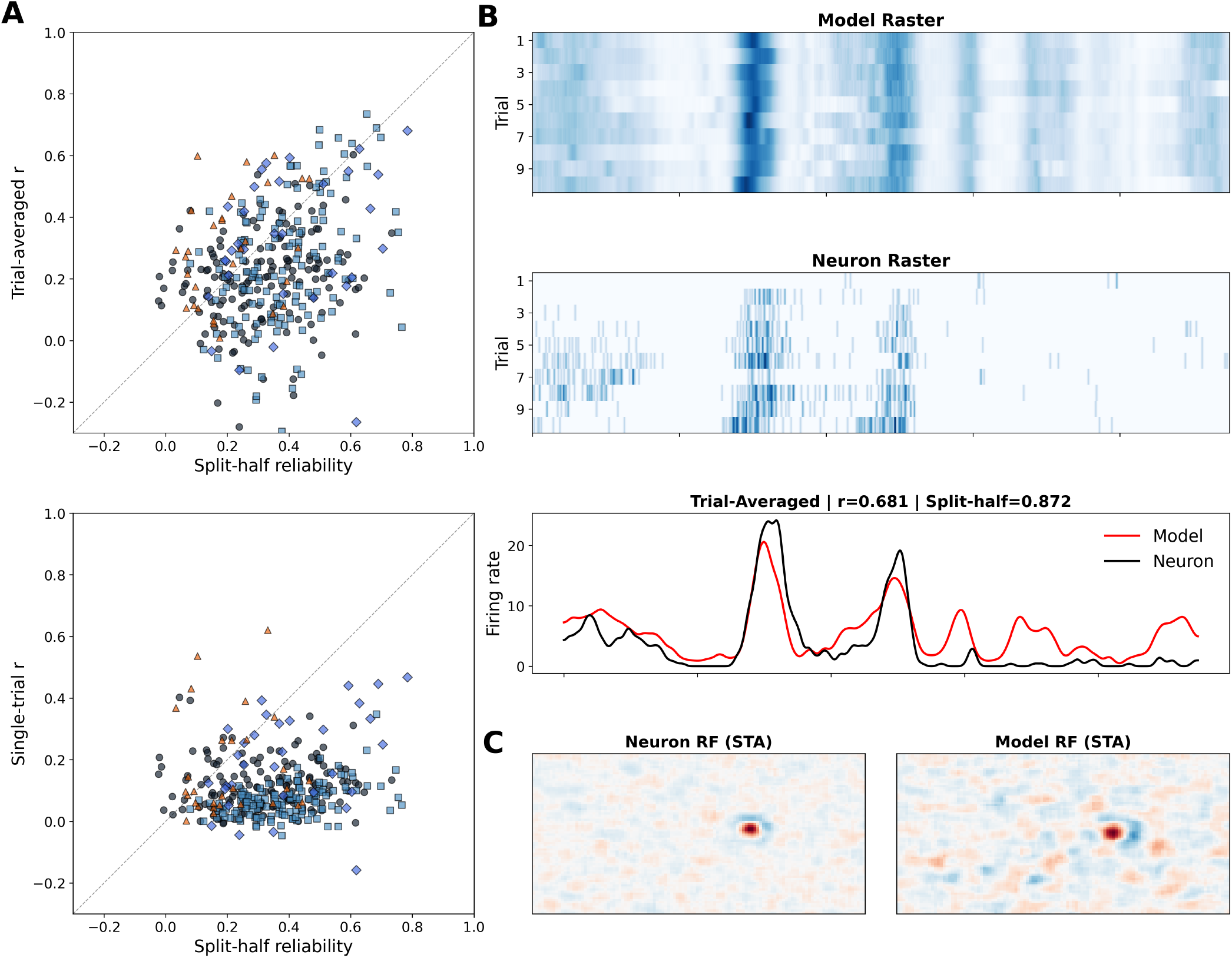
Model evaluation metrics for cortical V1 extracellular single-cell recordings. **(A)** Trial-averaged and single-trial prediction performance plotted against neural split-half reliability, for cortical CNN models across four animals (n = 348 cells); different symbols indicate different animals. **(B)** Example model–neuron evaluation on a single trial and on the trial-averaged prediction for held-out natural-scene repeats (n = 10 trials). **(C)** Example neuron spike-triggered average (left) and model spike-triggered average (right), measured with white noise.

We then used the CNN models to explore properties object motion sensitivity (OMS), and to generate hypotheses for further testing. We probed models with two stimulus classes, jittering differential motion as previously presented to the retina, and drift differential motion which contains information about orientation and spatial frequency.

We presented matched differential-motion (DM) and global-motion (GM) grating stimuli: a center disk and surrounding region that moved independently (DM) or coherently (GM). We presented two classes of object motion stimuli to models of the retina and cortex, which were qualitatively present in the natural images experienced by the mouse in vivo. The first probed the problem of object detection, with a stimulus composed of fine jitter with the statistics of fixational eye movements, as previously been shown to identify object motion sensitive cells of the retina. The second probed the problem of discriminating objects from each other using spatiotemporal pattern information about the object. These stimuli used drifting motion, as would occur from optic flow of a background created by body motion in combination with larger object motion. We presented a large range of spatial frequencies and center–surround orientations, using the model to test a wider range of conditions each centered on a unit’s receptive field, a set of stimuli that would have been practical in an experiment. For every unit we computed an OMS index as the difference in predicted response variability between DM and GM, normalized by total activity and averaged over conditions.

The two models revealed a functional double dissociation between the retinal and cortical OMS computation in the models (Fig. 3). The retinal model responded preferentially to jitter DM over drift DM, whereas the cortical model showed the opposite preference (Fig. 3 B,C). Moreover, cells showing object motion sensitivity in the cortex exhibited orientation selectivity, thus conveying specific information about the object pattern (Fig. S5-6). The complementary nature of these two strategies suggests that the cortex recomputes object motion sensitivity rather than directly inheriting it from the retina. To test this model-driven hypothesis, we conducted further experiments in mouse V1 and retina (Fig S7, S9). We presented the same matched pairs of DM and GM stimuli at two spatial frequencies, centering them on receptive fields localized prior to presenting the grating stimuli. As predicted by the model, retinal neurons showed greater preference for fine jitter DM than drift DM, whereas V1 neurons showed greater preference for drift DM than jitter DM. Also as predicted, V1 cells exhibiting object motion sensitivity were orientation selective (Fig. 3B, Fig. S8, S10). Given that oriented surround suppression and figure ground motion are known phenomenology in mouse V1 (25, 26), we asked whether orientation-gated OMS was related to oriented-tuned surround suppression (OTSS). However, the property of OTSS was only weakly correlated with object motion sensitivity, (Fig. S5-6, Fig. S8), indicating that the cortical orientation-gated OMS computation is not explained by orientation-tuned surround suppression alone.

**Fig. 3.**
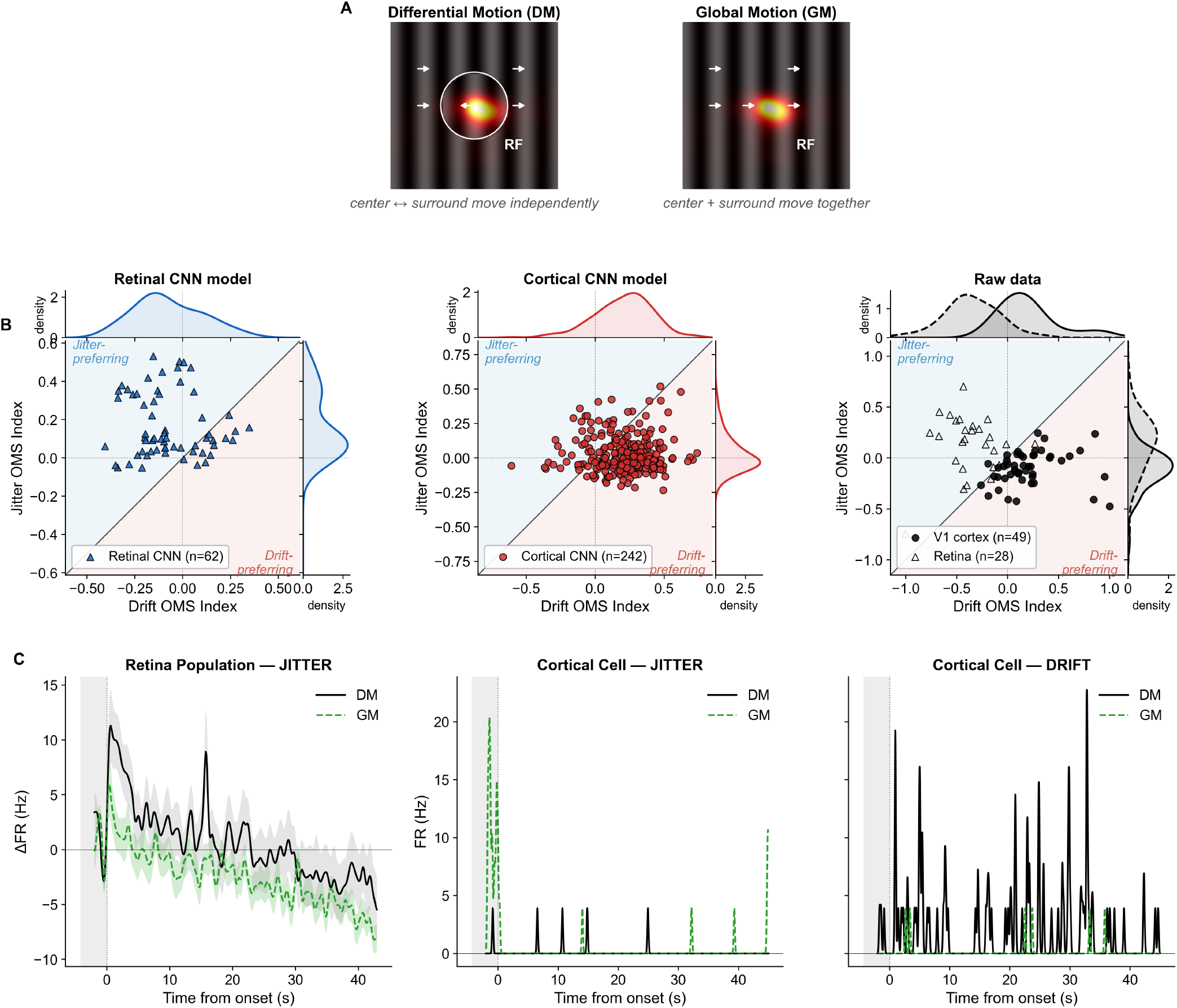
CNN models show the retina and cortex have distinct OMS computations. **(A)** Differential Motion (DM) vs. Global Motion (GM) stimulus paradigm. Center and surround move independently in DM; move together in GM. RF gradient overlay shows the receptive field of a representative cortical neuron. **(B)** OMS index (Drift vs. Jitter) scatter distributions across neural populations. Left: retinal CNN model (2 models-datasets, n=62 cells). Middle: cortical CNN model (4 models-datasets, n=242 cells). Right: raw biological data in targeted experiment for a separate mouse, V1 cortex (n=49 cells, filled circles) and a targeted OMS recording in a separate retina (n=28 cells, open triangles), max-abs normalized within the population. Jitter and Drift axis distributions from the model and recorded cells in the marginals. **(C)** Example recorded neural responses to jitter and drift stimuli. Left: retina population average (n=28 cells, p=0.025) showing DM (black) and GM (green) firing-rate responses. Middle and right: single V1 cell responses to drift and jitter differential- and global-motion stimuli. Retina and cortex OMS-index distributions differ significantly on the drift and jitter axes, in both the CNN and the recorded data (two-sample Kolmogorov-Smirnov; models: Drift D=0.55, Jitter D=0.45; data: Drift=0.80, Jitter D = 0.58; all *p* < 10^−4^), showing cortex more drift-preferring and retina more jitter-preferring)

If the computation of object motion sensitivity were completed in the retina and inherited by V1, one would expect that a linear readout could reproduce cortical object motion sensitivity (27). We tested this idea by freezing a retina-trained CNN model and then trained a linear map of its output to V1 responses (Fig. 4A). We additionally tested a frozen retina followed by a 3 layer CNN to predict V1 OMS responses, in case additional nonlinear processing were required. Both models predicted V1 OMS responses less well than V1 models trained to natural stimuli (fig. S4), and, critically, the cortical drift preference seen in both the end-to-end model and the in vivo population collapsed back toward the retinal jitter preference (Fig. 4B; fig. S3). No linear transformation of the retinal computation recovered the cortical one, and even three added nonlinear layers only partially reconstructed it, cortical drift-OMS was diminished whenever the model was constrained to build on retinal output. Even though our retina model could reproduce retinal properties of object motion sensitivity, it was insufficient to allow cortical object motion sensitivity to be reconstructed. This implies that cortical object motion sensitivity is constructed in part from retinal populations or properties other not included in our retinal model, beyond those that create retinal object motion sensitivity.

**Fig. 4.**
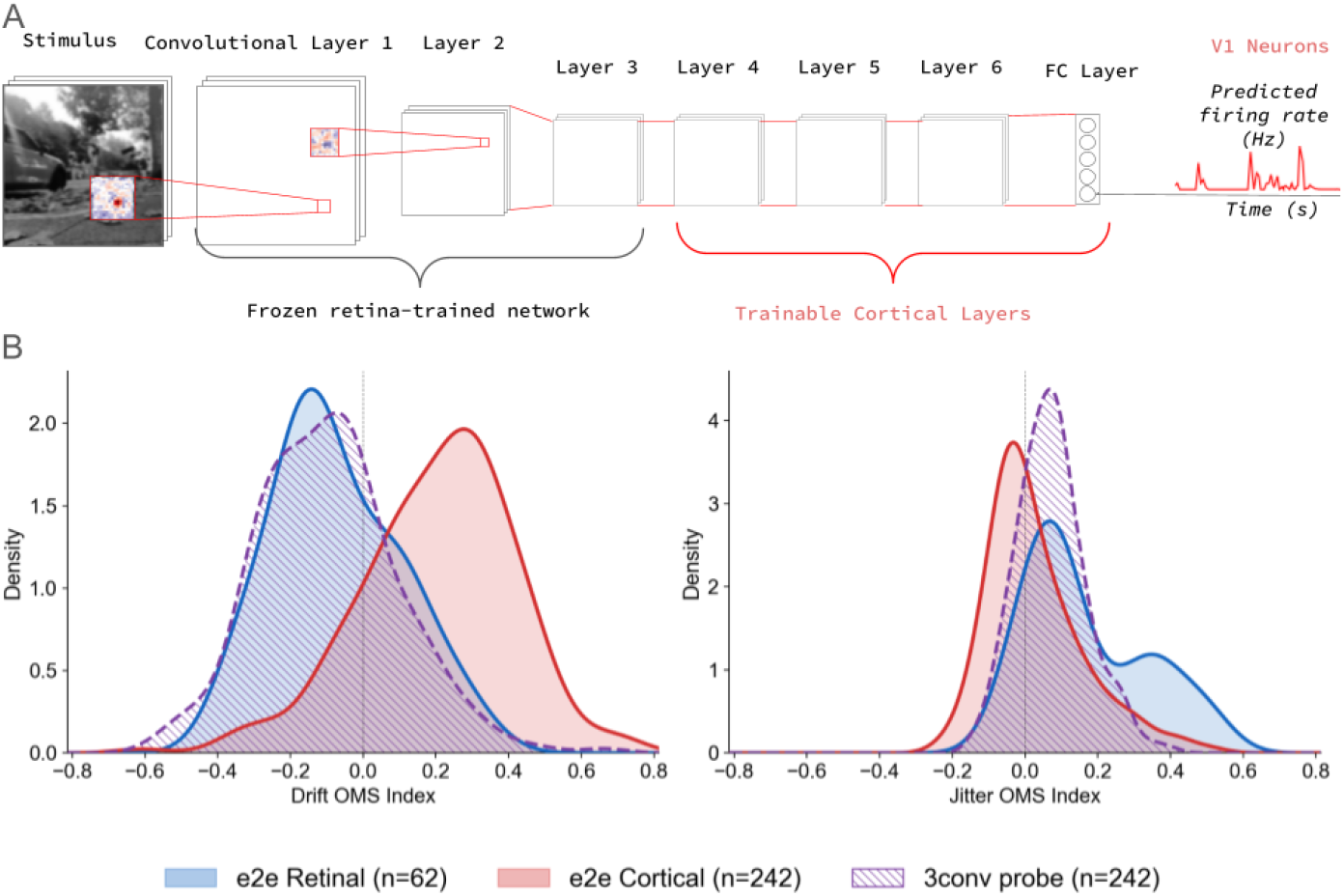
Retino-cortical probe testing whether cortical object motion sensitivity is a readout of retinal output. **(A)** Schematic of the retino-cortical probe. A convolutional network trained end-to-end (e2e) to predict retinal responses is frozen and followed by a linear readout, or by three trainable convolutional layers, fit to predict V1 firing rates to natural scenes across all four cortical datasets. **(B)** Distribution of the OMS index in response to global or differential motion for the end-to-end retina (blue), end-to-end cortex (red), and the retino-cortical probe (purple striped), on the drift and jitter axes. End-to-end retinal units are jitter-preferring and drift-suppressed (drift median = −0.10, jitter median = +0.11), whereas cortical units are drift-preferring and jitter-suppressed (drift median = +0.23, jitter median = +0.00); Kolmogorov–Smirnov: drift D = 0.55, *p* = 2 × 10^−14^; jitter D = 0.45, *p* = 1 × 10^−9^. The retino-cortical probe is statistically indistinguishable from the retinal jitter response (KS D = 0.11, *p* = 0.57, Mann– Whitney *p* = 0.45) and remains strongly separated from the end-to-end cortical drift response (D = 0.61, Mann–Whitney *p* = 6 × 10^−42^).

To further examine whether cortical object motion sensitivity was inherited from the retina, using the same frozen-retina 3-CNN layer model, we computed the contribution of each model retinal ganglion cell type to each cortical neurons using the method of Integrated Gradients (24, 28, 29). This gave the weight of each ganglion cell type to each cortical cell type for a specific DM or GM stimulus. From these contribution weights, we computed the contribution-weighted average of the OMS indices of the retinal channels to determine the cortical OMS behavior could be predicted from the weights of the retinal cells (Fig. S11). If V1 inherited OMS from dedicated OMS-selective channels, this prediction would track the measured cortical OMSI. However, we found this was not the case. Sensitivity of a cortical cell to drift differential motion was not correlated with that of the retinal cells that contributed to that cortical neuron. Jitter selectivity, in contrast, showed a weak correlation that was carried by the inhibitory contributions of the retinal cells to the cortical cell (Table S1). This indicated that object motion sensitivity for drifting stimuli was not directly inherited from properties of retinal neurons, but rather reconstructed after the level of the retina.

As an illustration of how populations of OMS cells respond to natural motion, we presented to population models of retinal and cortical OMS cells a movie from the Moving Camouflaged Animals (MoCA) dataset (30). In an example of an arctic fox (Figure 5; Video S1). We find that the retinal activation heatmaps are more focal to animal motion, from crouching to jumping, showing a more spatially invariant response to object motion. Cortical model activations, in contrast, carried information about the animal orientation and produced an extended activation pattern that spanned the entire animal’s body (Figure 5B, Fig S12). We decoded the population orientation and direction vectors by linearly weighting unit activity with their normalized tuning curves, revealing that while per-cell orientation selectivity similar across the network, the cortical OMS population exhibits stronger direction selectivity and responds to a broader range of orientations than non-OMS cells (Video S2). Together, this suggests a division of labor in which the retina flags that something is moving while the cortex resolves where the moving body is and how it is oriented, which adds spatial specificity an animal would need to detect and orient themselves with respect to a moving object such as a predator.

**Fig. 5.**
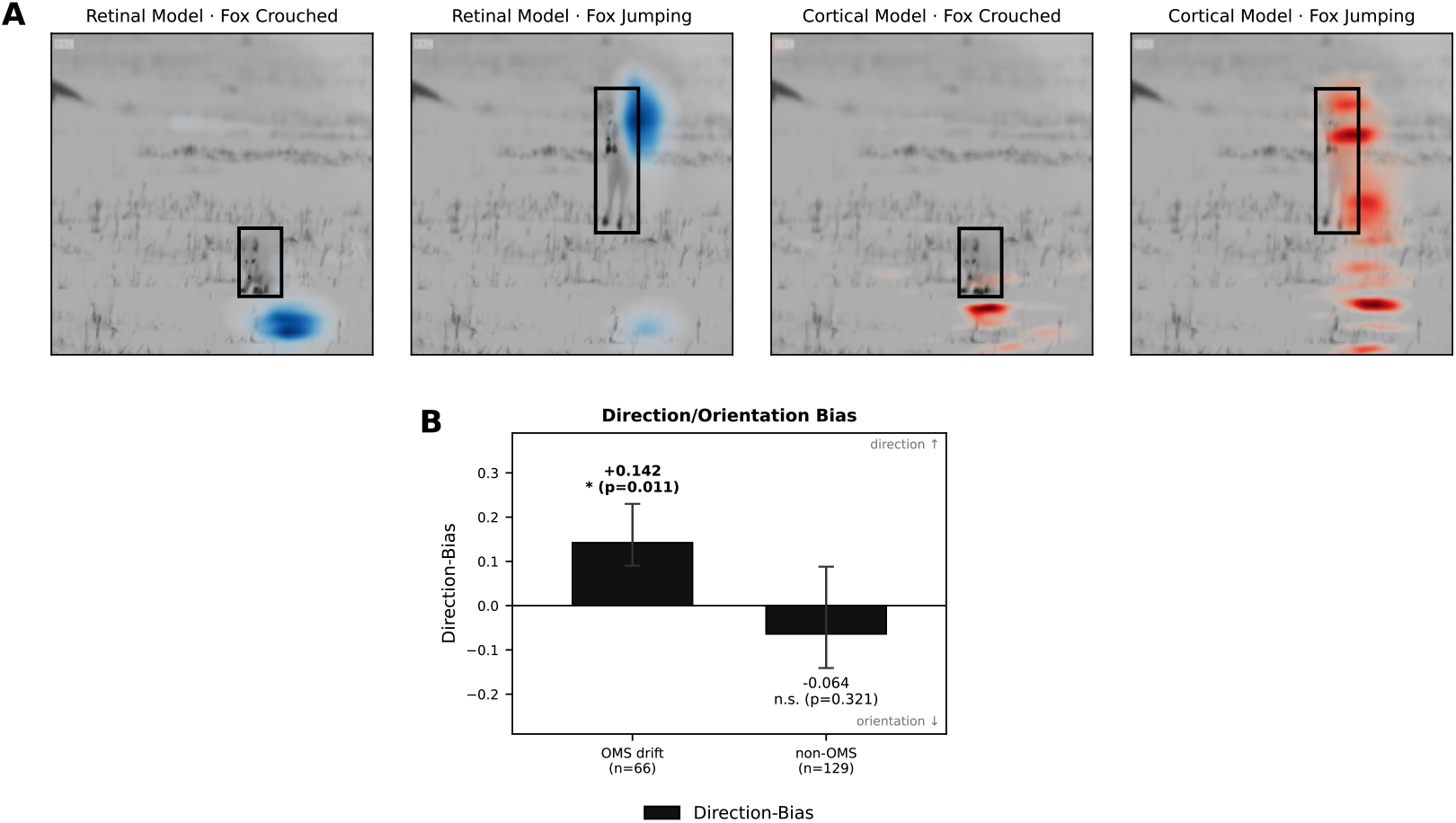
Cortical activation concentrates on the moving animal and is more spatially structured than retinal activation. **(A)** Model activation (retinal, blue; cortical, red) overlaid on the raw frame for a crouched and a jumping frame from an arctic fox of the MoCA dataset. The black box is the ground-truth annotation of the animal’s location and containment. **(B)** Per-cell Direction-Bias = (DSI − OSI)/(DSI + OSI) for the cortical model’s independently identified object-motion-sensitive units (OMS drift, n = 66) versus the remaining units (non-OMS, n = 129), from full-field drifting gratings.

## Discussion

Here we have used matched stimuli in the retina and in primary visual cortex to compare in detail processing strategies to represent object motion against a background. The models we have used to describe and identify properties of object motion sensitivity were derived from natural stimuli containing differential motion, indicating the coding properties we have described are relevant to processing of visual stimuli. The use of models as an intermediate was also essential to exploration of a wide range of stimuli in order to develop efficient experimental tests of those coding properties. The functional specialization we identify in retina and cortex fits with likely ethological functions. Retinal cells prefer fine jitter differential motion, which is useful for detecting the smallest deviation of object motion from background motion due to incessant fixational eye movements. The price retinal cells pay for such sensitivity is to sacrifice information about the specific visual pattern (1, 31). In contrast, cortical object motion sensitivity conveys information about the specific spatial pattern. Seen this way, it makes sense as to why the cortical computation would not directly inherit output from retinal OMS cells, as they had discarded some spatial pattern information. In the transmission to higher levels of the brain, one would expect a conjunction of the signals from retinal OMS cells specialized in object detection with cortical OMS cells that encode pattern information, with the resulting representation consisting of both sensitive detection, and information about object identity.

The visual hierarchy is commonly understood as a sequential processing pipeline of progressive refinement. The standard expectation (9–11) is that V1 elaborates upstream computations. With respect to object motion, this would suggest inheritance of a single computation for purposes of efficiency. In the case of object motion, the needs of the task make inheritance a suboptimal solution, whereas parallel specialization and recomputation at different levels brings advantages (32, 33). One must consider that other computations solved at one level may likewise be rebuilt downstream for different purposes.

## ACKNOWLEDGEMENTS

The authors would like to thank Bob Schneeveis, Tom Clandinin, and Krishna Shenoy for their valuable contributions. We thank Steven W. Zucker and Luciano Dyballa for helpful discussions on the application of their manifold method to our retinal and cortical ANN models.

## AUTHOR CONTRIBUTIONS

Conceptualization: JCW, SAB Methodology: JCW, DA, JBM, YF, SAB Investigation: JCW, SAB, DA, YF, ZA Visualization: JCW, DA, ZA Funding acquisition: SAB Project administration: SAB Supervision: SAB Writing – original draft: JCW, SAB Writing – review & editing: JCW

## Methods and analysis

### Animal subjects

All procedures were approved by the Stanford University Administrative Panel on Laboratory Animal Care (APLAC protocol #11619) and conformed to NIH guidelines. Adult male C57BL/6J mice were obtained from the Jackson Laboratory and aged 8–22 weeks at the time of recording. Mice were entrained to a 12 h light/dark cycle (lights on 07:00, off 19:00) and recordings were performed at dawn and dusk (05:00 or 18:00).

### Surgery

Surgeries were performed under aseptic technique with the mouse held in a stereotaxic carrier on a self-regulating heating pad, under isoflurane anesthesia (0.5–3%) with ophthalmic ointment applied to prevent corneal drying and subcutaneous warmed saline (0.9% NaCl) given through the procedure; anesthetic depth was monitored by breathing rate, body temperature, toe-pinch response, and mucous-membrane color. Pre-operative analgesia was provided with buprenorphine SR (0.5–1 mg/kg s.c.) or Ethiqa XR (extended-release buprenorphine, 3.25 mg/kg s.c.), and carprofen (5–10 mg/kg s.c.) was given at the end of surgery; supplemental buprenorphine HCl (0.05 mg/kg s.c., every 6–12 h) was administered if signs of pain were observed.

In a first surgery, the skull was exposed through a midline incision and a head plate (8 mm diameter) was attached with a dental cement/acrylic mix to permit head fixation; a gold reference pin was implanted over frontal cortex to serve as the recording reference together with the probe tip. Mice recovered for 2–3 days before habituation. Approximately two to four weeks later, in a second surgery, a small craniotomy (∼0.5 mm diameter) was opened over primary visual cortex, its location determined from a stereotaxic atlas and skull landmarks and drilled superficially enough to expose the dura while the skull was kept moist with sterile saline. Between sessions the craniotomy was sealed with the silicone elastomer Kwik-Sil and protected by a headplate cover. Immediately before each recording the Kwik-Sil was removed, agar was applied over the craniotomy, and a drop of silicone oil was placed on the agar to prevent brain tissue dehydration.

### Habituation and head fixation

Beginning approximately six days before recording, mice were habituated to handling and head fixation on an escalating schedule. On the first day mice received gentle handling (20 min) and 5 min on the rig in the light with no stimulus and no head fixation; on the second and third days, gentle handling (20 min) followed by 30 and 45 min of head-fixed rig time in the light with no stimulus. On the fourth day mice received 30 min on the rig in the light, then 30 min in darkness with white noise, then 30 min in darkness with natural scenes; the fifth day repeated this structure with longer blocks (up to 60 min each) and, for some animals, a final shorter natural-scene block or a delayed (ZT10) session. Mice ran freely on a cylindrical treadmill throughout.

### In vivo electrophysiology

Acute extracellular recordings were made from V1 with Neuropixels probes — a four-shank Neuropixels 2.0 probe in most sessions and a Neuropixels 1.0 probe in others (35). With the four-shank probe the shanks were inserted at a 30° angle so that all four shanks remained normal to the cortical layers. The reference was the probe tip, in some sessions combined with the frontal gold pin. The probe was advanced under micromanipulator control to a depth of interest (for example, ∼1.1–1.4 mm). Receptive fields were confirmed online before recording using a sparse-noise mapping stimulus and confirmed using spike amplification through SGLX computer speakers.

Neural data were acquired in SpikeGLX (IMEC stream). A multi-channel data-acquisition system (NIDAQ) recorded the photodiode signal from the stimulus monitor, the rotary-wheel locomotion signal, and the camera strobe; a second acquisition stream (LabJack, ∼5 kHz) recorded the strobe, photodiode (stimulus and square-wave channels), wheel channels, and the Neuropixels start and sync TTLs. Recording onset was verified by monitoring the Neuropixels-start channel for the transition coincident with SpikeGLX streaming. The IMEC/NIDAQ, Jackfish, and StimPack clocks were aligned post hoc: StimPack per-frame timestamps were anchored to the NIDAQ and Jackfish photodiode signals, and residual clock drift between streams was measured at paired temporal anchors and removed by linear interpolation onto a common time base. Continuous behavioral signals were resampled to this common base by linear interpolation, and spike counts by cumulative-sum interpolation followed by differentiation.

Spikes were sorted with Kilosort 4 (34). Raw IMEC data were preprocessed with a 300–8000 Hz bandpass filter, common-median referencing across channels, and phase-shift correction; bad channels were detected and removed, and drift/motion correction was performed within the Kilosort 4 pipeline. Sorted units were curated in Phy, and only units labeled “good” were retained. For each retained unit the mean waveform was computed from 100 spikes over an 83-sample window (41 samples before and after the spike trough; 30 kHz sampling) on the unit’s peak-amplitude channel, and refractory-period quality was assessed from autocorrelograms for the inter-spike-interval (1 ms bins, ±50 ms window). Firing rates were estimated by binning spike times and convolving with a normalized Gaussian kernel; bin width and kernel standard deviation were matched per recording. We assessed cell stability by single-trail reliability at the start and then of the stimulus white noise and natural scene battery. Lastly, we factored cells into their respective linear receptive field response properties using the spike triggered average with respect to white noise stimuli shown during the recording.

After recording, probes coated with a fluorescent dye (per-animal dye color logged) were used for track reconstruction in animals designated for histology.Probe tracks were localized to V1 by reference to a standard atlas (Paxinos and Franklin, The Mouse Brain in Stereotaxic Coordinates, Elsevier, 2001).

### Visual stimulation

Stimuli were generated and presented with StimPack (https://github.com/ClandininLab/stimpack), an open-source client–server stimulus framework with a perspective-corrected rendering pipeline. Stimuli were displayed on a 24-inch 1920 × 1080 IPS monitor at 165 Hz, positioned 24 cm from the eye, with a photodiode mounted on the monitor for per-frame timing; analog signals from the photodiode, locomotion wheel, and camera strobe were acquired through a multi-channel data-acquisition system (36). The screen irradiance at the eye was 14.2–50.4 µW/cm^2^ during active stimulus presentation, 0.59 µW/cm^2^ during inter-stimulus periods, and 0.25 µW/cm^2^ during dark intervals (36). Stimuli were presented open-loop (independent of behavior).

Each session followed a fixed nine-block structure. Three repeated (test) blocks of repeated white noise, repeated natural-scene movie 1, and repeated natural-scene movie 2 were presented at the start of the session, followed by three unique (training) blocks of white noise and two natural-scene movies, and the three repeated test blocks were repeated at the end of the session, bracketing the training data. White-noise stimuli were spatially binned random checkerboards regenerated offline from stored random seeds. Natural-scene stimuli were hour-long continuous movies filmed at rodent eye level with an omnidirectional camera, spanning optical flow, head movements, translations, rotations, sudden stops, and still periods. For each frame, the stimulus reaching the retina was reconstructed by applying the measured gaze-dependent pixel shift (see Eye tracking); this gaze-corrected stimulus was the input to all encoding models, and the uncorrected screen content was retained for control analyses.

### Eye tracking and gaze-corrected stimulus reconstruction

Eye movements were tracked with a single-camera, calibration-free corneal-reflection system (36). An infrared-sensitive camera (Teledyne FLIR Grasshopper3, up to 90 fps, with a zoom lens) imaged the eye under 850 nm infrared flood lamp illumination, and two or more stationary fiducial infrared LEDs (850 nm, chosen not to interfere with visual stimulation) at known three-dimensional positions produced the corneal reflections used to set the angular scale. In each frame the pupil and the LED and camera corneal reflections were localized by a difference-of-Gaussians pipeline: the contrast-normalized, inverted image was raised to a tunable exponent, filtered by a difference of two Gaussians, thresholded, and the connected component nearest a seed point was taken as the centroid, with independent parameters for the pupil and fiducial pipelines.

Gaze was recovered in calibrated angular units by a geometric model using three coordinate systems: a world frame specifying the three-dimensional positions of the eye, camera, and each LED; the camera-pixel image frame; and an eye-centered spherical frame whose polar axis is the eye-to-camera line, with angular position given as elevation and azimuth. World positions of the camera and LEDs were recentered on the eye and rotated into the eye basis to yield static eye-centered LED angles; per frame, the pixel displacements of the pupil relative to the camera and fiducial reflections were converted to eye-centered elevation and azimuth through these LED angles. The model requires no per-animal eye-geometry parameter. A single residual offset of the camera reflection was determined by minimizing the disagreement among the per-LED gaze estimates at a reference frame in which all LEDs were visible. The system was validated to sub-degree accuracy against a rotary-encoder-controlled artificial eye over the ±20° range of mouse eye movements. Recording geometries (eye, camera, and fiducial-LED coordinates) were stored per recording period.

Blinks were detected from abrupt frame-to-frame jumps in the tracked coordinates and from pupil-coordinate outliers, with detected runs constrained to a plausible duration; blink frames were flagged, and the stimulus was blacked out during blinks for model input. The eye-centered gaze was converted to a per-frame pixel shift of the displayed image to produce the gaze-corrected stimulus, an estimate of the image on the retina given eye position. Gaze speed for eye-movement analyses was computed as the central-difference angular velocity on the sphere with the cosine-of-elevation azimuth correction, and locomotion was converted from encoder ticks to cm/s estimated by a 30.48 cm wheel and 1024 ticks per revolution.

### Ex vivo retinal electrophysiology

Mice were dark-adapted for at least three hours (up to overnight) before the experiment, and all tissue preparation was performed under dim red light. Mice were anesthetized with isoflurane and euthanized by cervical dislocation, after which both eyes were enucleated and hemisected along the cornea. The eyecup was dissected at room temperature in Ames’ medium (Sigma-Aldrich A1420) buffered with sodium bicarbonate and equilibrated with 95% O_2_ / 5% CO_2_, under infrared (*>*850 nm) illumination through a stereomicroscope with night-vision optics, and the vitreous and pigment epithelium were removed to isolate the retina (as previously described; Lee, Kim, and Baccus, bioRxiv 2024.03.18.585364, 2024). The isolated retina was mounted ganglion-cell-side up on a multi-electrode array and held in place by, and was continuously superfused with oxygenated, bicarbonate-buffered Ames’ medium held at 30–32 °C by a temperature-controlled stage. The retina was allowed to rest on the MEA for at least an hour before recording. Retinal ganglion cells were recorded extracellularly on a sampling rate of 30 Khz, and spikes were sorted with an in-house spike sorting software under the same unit-quality criteria as the in vivo recordings. The same gaze-shifted natural-scene videos presented to the mice in vivo were projected onto the photoreceptor layer.

### Encoding models

Retinal and cortical responses were predicted from the gaze-corrected stimulus by convolutional neural networks fit with the same framework, each taking a temporal stimulus window as input and reading out non-negative predicted firing rates through a final Softplus nonlinearity. Convolutional layers used non-affine LayerNorm, a Softplus nonlinearity, and dropout (*p* = 0.1). The cortical model (e.g. goldroger) took a 36-frame window and comprised four to six convolutional layers; a first layer of 14 filters of size 16 × 16 and three further 14-channel layers of size 14 × 14, 12 × 12, and 8 × 8 then a fully convolutional readout layer (8 × 8 filters) producing one feature map per cell, and a spatial readout over the 19 × 19 output grid, predicting 195 units. The retinal model took a 30-frame window and comprised two or three convolutional layers of 16 channels (first-layer filter 30 × 30, subsequent filters 22 × 22) and a flattened fully connected readout to the recorded ganglion cells; the two retinal models read out 36 and 26 cells.

Models were trained to minimize the Poisson negative log-likelihood between predicted and measured firing rate, with *L*_1_ penalties on the unit activations and on the network weights using the AdamW optimizer. The stimulus was normalized before input and frames during blinks were zeroed. Stimulus and responses were resampled to a common model frame rate of 50 Hz. The unique blocks were used for training and the repeated blocks for evaluation; inputs were sliding temporal windows preceding each response time.

Let *λ*_i,t_ = *f*_ϕ_(*s*_t_)_i_ be the model’s predicted rate for unit *i* at frame *t* (the network output after the final Softplus, so *λ*_i,t_ *>* 0), and *y*_i,t_ the measured rate. Models minimize the Poisson negative log-likelihood plus two *L*_1_ penalties:

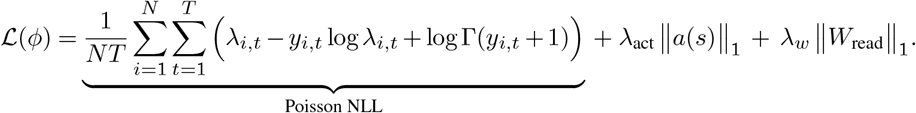

- The log Γ(*y* + 1) term is the full (“Stirling”) normalizer, included so the loss is a proper NLL (`PoissonNLLLoss(log_input=False, full=True)`).
- ∥*a*(*s*)∥_1_ is the *L*_1_ penalty on convolutional-unit activations (activity sparsity); ∥*W*_read_∥_1_ = ∑|*W*_read_| is the *L*_1_ penalty on the readout weights, with *λ*_w_ = 10^−5^.
- Optimizer AdamW; inputs contrast-normalized; blink frames zeroed; the unique blocks train, the repeated blocks evaluate.

Each convolutional block applies non-affine LayerNorm, Softplus, and dropout (*p* = 0.1):

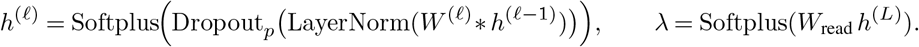

Modeling was restricted to a curated set of units per recording (∼24–214 neurons per animal), further filtered by held-out predictive correlation and split-half reliability. For receptive-field-centered analyses, the gaze-corrected stimulus was resampled around the unit’s receptive-field center by a differentiable grid sampler (bilinear interpolation, zero padding) to a fixed crop.

### Model evaluation

Model quality was assessed with three measures. Split-half reliability was the Pearson correlation between firing-rate estimates from two independent halves of the repeated trials, computed over 30 random partitions. Trial-averaged correlation was the Spearman correlation between the mean model prediction and the mean measured response across repeats. Single-trial correlation was the Spearman correlation against each individual trial using that trial’s gaze-shifted stimulus. Model receptive fields were computed as the gradient of the unit’s output with respect to the stimulus and spike-triggered averages by reverse correlation over the stimulus-history window; both were decomposed into separable spatial and temporal components by truncated singular-value decomposition of the spatiotemporal kernel and compared in position, structure, and orientation to the measured receptive field on white-noise stimuli.

### Object-motion stimuli and object-motion index

Object motion was probed with matched differential-motion and global-motion grating stimuli (after 1), each comprising a central disk and a surround filled with a drifting or jittering sinusoidal grating. In the differential-motion condition the center and surround moved independently (opposite drift directions, or independent jitter trajectories); in the global-motion condition the entire field moved coherently. In drift conditions the center and surround gratings drifted in opposite directions at a fixed temporal frequency; in jitter conditions each followed an independent random walk about its mean position. Stimuli spanned all sixteen combinations of center and surround orientation from 0°, 45°, 90°, 135° and were centered on each unit’s receptive field. For the trained models, the stimulus used spatial frequencies of 0.04–0.08 cycles per pixel, a 20-pixel center-disk radius, and a jitter random walk of up to 15 pixels per frame, was centered on the unit’s receptive field determined by gradient-based reverse correlation, and was preceded by a gray lead-in period excluded from response statistics; spatial frequencies, disk size, and centering for the targeted recordings are given below.

#### Model OMSI (per unit)

With *σ*_•_ = std_t_ [*λ*_t_] the temporal standard deviation of the predicted rate over the stimulus period (drop the gray lead-in), computed separately for the DM and GM conditions:

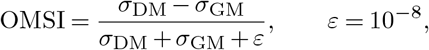

evaluated once with coherent drift (OMSI_drift_) and once with random-walk jitter (OMSI_jitter_).

#### Measured OMSI (per recorded unit)

*σ* is replaced by the mean firing rate 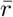 over each matched trial segment, taken per DM/GM pair, medianed across pairs within a dynamic condition:

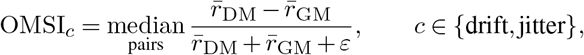

and the reported combined index is the mean of the two conditions,

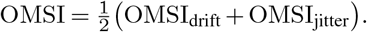

Units whose DM or GM segment rate falls below a low-activity threshold (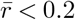 spikes s^−1^) are excluded from that pair.

**Orientation selectivity** (drifting gratings at *M* = 8 orientations *θ*_m_, response *r*(*θ*_m_)):

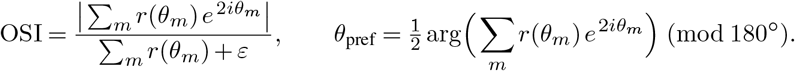

The double angle 2*θ* makes the index 180°-periodic; the direction selectivity index (DSI) is the identical construction with *e*^iθ^ (single angle, 360°-periodic) and *θ*_pref_ = arg(·) (mod 360°).

Differences in receptive-field structure between object-motion-sensitive and non-object-motion-sensitive populations were assessed by bootstrap: cells in each pool were resampled with replacement over 1000 replicates, and the per-pixel difference between pool means was reported as significant only where the 95% bootstrap confidence interval excluded zero.

### Targeted object-motion experiment

A targeted object-motion experiment was run in mouse V1 (Neuropixels 1.0) and, separately, in the ex vivo retina on the multi-electrode array, using a single fixed stimulus sequence (Supplementary Figs. 6 and 7). The sequence comprised a white-noise block for receptive-field mapping; a full-field sweep of drifting gratings at eight orientations (0–315° in 45° steps; OS_Beginning blocks, 1800 frames ≈ 36 s each at the 30 Hz stimulus rate); continuous differential- and global-motion jitter blocks (9000 frames ≈ 180 s); a battery of 128 center–surround grating blocks (1350 frames ≈ 27 s each, 0.6 s blank inter-stimulus intervals); and a repeat of the eight-orientation sweep at the end (OS_End blocks). The 128 grating blocks fully crossed motion type (64 differential, 64 global), temporal dynamics (64 drift, 64 jitter), spatial frequency (0.02 and 0.04 cycles per pixel), and all sixteen center–surround orientation combinations from 0°, 45°, 90°, 135° ; checkerboard differential/global jitter blocks were also included.

Receptive-field centers were assigned from the white-noise block by a rate-weighted spatiotemporal spike-triggered average over the preceding 8 frames, taking the spatial receptive field at the lag of peak energy and the intensity-weighted centroid of the region exceeding three times a robust (median-absolute-deviation) noise estimate. Only cells whose centroid fell within the central differential-motion disk were included (n = 51 of 237 sorted units, V1; n = 28 of 44 cells, retina; Supplementary Figs. 6 and 7). Orientation tuning was computed from the eight oriented-grating blocks, averaging the beginning and end sweeps and discarding the first 150 frames of each block. Firing-rate object-motion indices (drift and jitter) were computed as above; retinal firing rates were taken at 20 ms resolution.

### Retinal-cortical probes and integrated-gradient inheritance test

Two probe models sharing a frozen, retina-trained convolutional front end were fit to the V1 population. The linear probe applied the frozen retinal fully convolutional trunk followed by a single trainable linear layer. The three-convolution probe applied three randomly initialized trainable convolutional layers, each with a nonlinearity, and a fully connected readout, trained under the Poisson loss used for the end to end encoding models. Both probes were compared to the cortical model trained end to end on V1 data by single-trial held-out Pearson correlation. Per-channel retinal contributions to each cortical cell were quantified by integrated gradients (28) on the three-convolution probe, whose frozen retinal front end contained 26 output channels.

#### Attribution

For cortical output *F*_c_ (cell *c*), retinal front-end channel *k*, zero-stimulus baseline *x*^′^ = 0, and *m* path steps (*m* = 5), integrated gradients are accumulated by the trapezoidal rule along the straight-line path 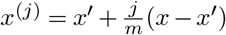:

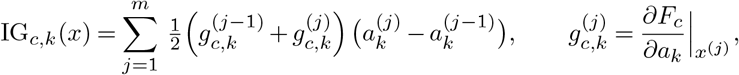

where 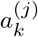 is the channel-*k* activation (summed over space, *H* × *W* ) at path point *j* and 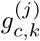 the corresponding gradient. (This trapezoidal path integral *g* d*a* is the implemented form; it reduces to the textbook 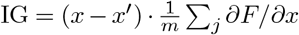 for a linear activation.)

**Excitatory/inhibitory split** by sign:

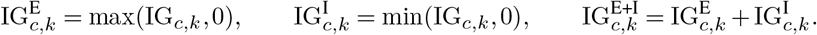

#### Inheritance test 1 — ΔContrib vs ΔFiring

Averaging each quantity over the *W* ≈ 921 windows of a condition, define per cortical cell *c* (population *n* = 33) and channel *k*:

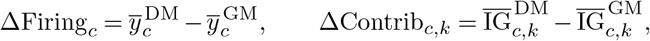

and report, per channel, the Pearson correlation across the 33 cells

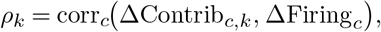

#### Inheritance test 2 — contribution OMSI

With 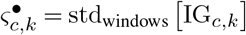 the temporal std of channel *k*’s contribution to cell *c*:

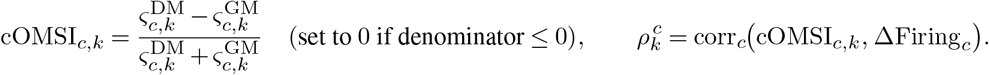

#### Predicted (inherited) cortical OMSI

A contribution-weighted average of the retinal channels’ own OMSI, with weights the absolute contribution magnitude of each channel to that cell (E, I, or E+I; Supplementary Table 1):

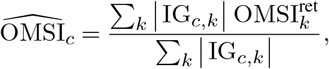

compared against the measured cortical OMSI_c_ by Pearson correlation with a shuffle null.

Model instantaneous receptive fields were estimated from unit-output gradients across stimulus windows and decomposed by singular-value decomposition via the Gram matrix, taking the top component as the linearized receptive field. Receptive-field centers were defined from its peak-lag energy, and a unit was retained when its natural-scene and white-noise instantaneous-receptive-field centers agreed to within 5 pixels.

### Population vector analysis

#### Unit Responses and Tuning

For each model unit *i* at spatial location (*x, y*), we define a direction-tuning vector, *F*_x,y,i_(*θ*). This vector represents the mean response to an object-motion-sensitive (OMS) differential-motion grating drifting in one of eight directions:

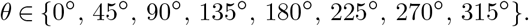

Each tuning vector is normalized to unit length. For a given video frame *t*, we extract the baseline-subtracted unit activations, denoted as *A*_x,y,i_(*t*).

#### Population Response

The population orientation and direction response for a given pixel and frame is calculated by projecting the instantaneous activity onto the unit’s tuning profile:

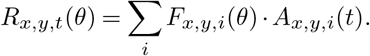

#### Circular Vector Summation

We apply a circular vector sum of *R*_x,y,t_(*θ*) over the eight directions to obtain the local vector fields.

##### Direction Vector (360°-periodic)

The direction vector components are computed as follows. Horizontal Component: *H*_dir_ = ∑_θ_ *R*(*θ*) cos(*θ*). Vertical Component: *V*_dir_ = ∑_θ_ *R*(*θ*) sin(*θ*). Magnitude: 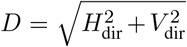. Heading (Angle): *φ*_dir_ = atan2(*V*_dir_, *H*_dir_).

##### Orientation Vector (180°-periodic, double-angle)

The orientation vector components are computed using a double-angle projection. Horizontal Component: *H*_ori_ = ∑_θ_ *R*(*θ*) cos(2*θ*). Vertical Component: *V*_ori_ = ∑_θ_ *R*(*θ*) sin(2*θ*). Magnitude: 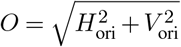 . Axis (Angle): 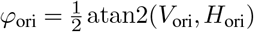.

### Instantaneous model receptive fields

The instantaneous receptive field of unit *i* for stimulus window *s* is the input gradient of that unit’s scalar output,

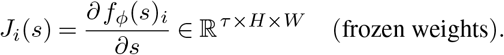

Stacking the *n* windowed gradients as rows of *X* ∈ ℝ^n×D^ (*D* = *τ HW* ), the linearized RF is the leading component of the *n* × *n* Gram matrix,

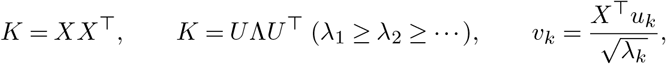

where 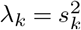 are the squared singular values of *X* and *v*_1_ (reshaped to *τ* × *H* × *W* ) is the top spatiotemporal component. It is separated into temporal and spatial parts by peak energy:

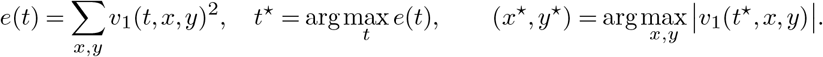

A unit is retained when its natural-scene and white-noise RF peaks agree,

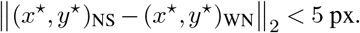

## Data and code availability

Stimuli were generated with StimPack (https://github.com/ClandininLab/stimpack). Eye tracking used the corneal-reflection system of Au, Melander et al. (36). Spike sorting and data processing used Kilosort 4 and the lab processing pipeline; encoding models, object-motion analyses, and integrated-gradients attribution used the lab modeling and analysis packages. Processed datasets are packaged as HDF5 files.

## Supplementary Information

**Fig. S1.**
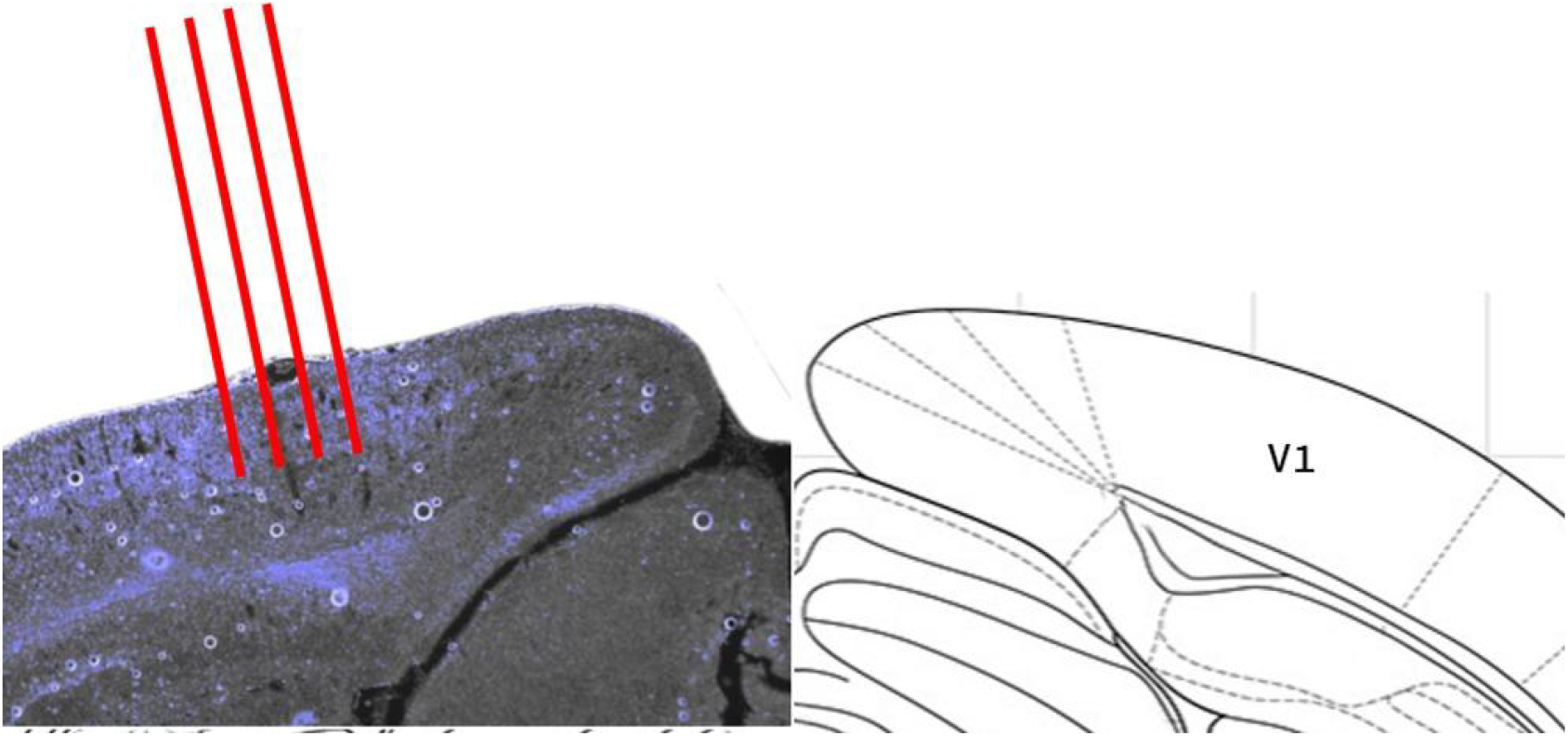
Histology from Neuropixel 2.0 recording in mouse V1 at a 30 degree angle to keep all four shanks normal to the cortical layers. Red lines indicating where the probes were inserted. Example histology from the Kizaru dataset in figure 1(referenced from Paxinos, George, and Keith B.J. Franklin. The mouse brain in stereotaxic coordinates: Access Online via Elsevier, 2001.)

**Fig. S2.**
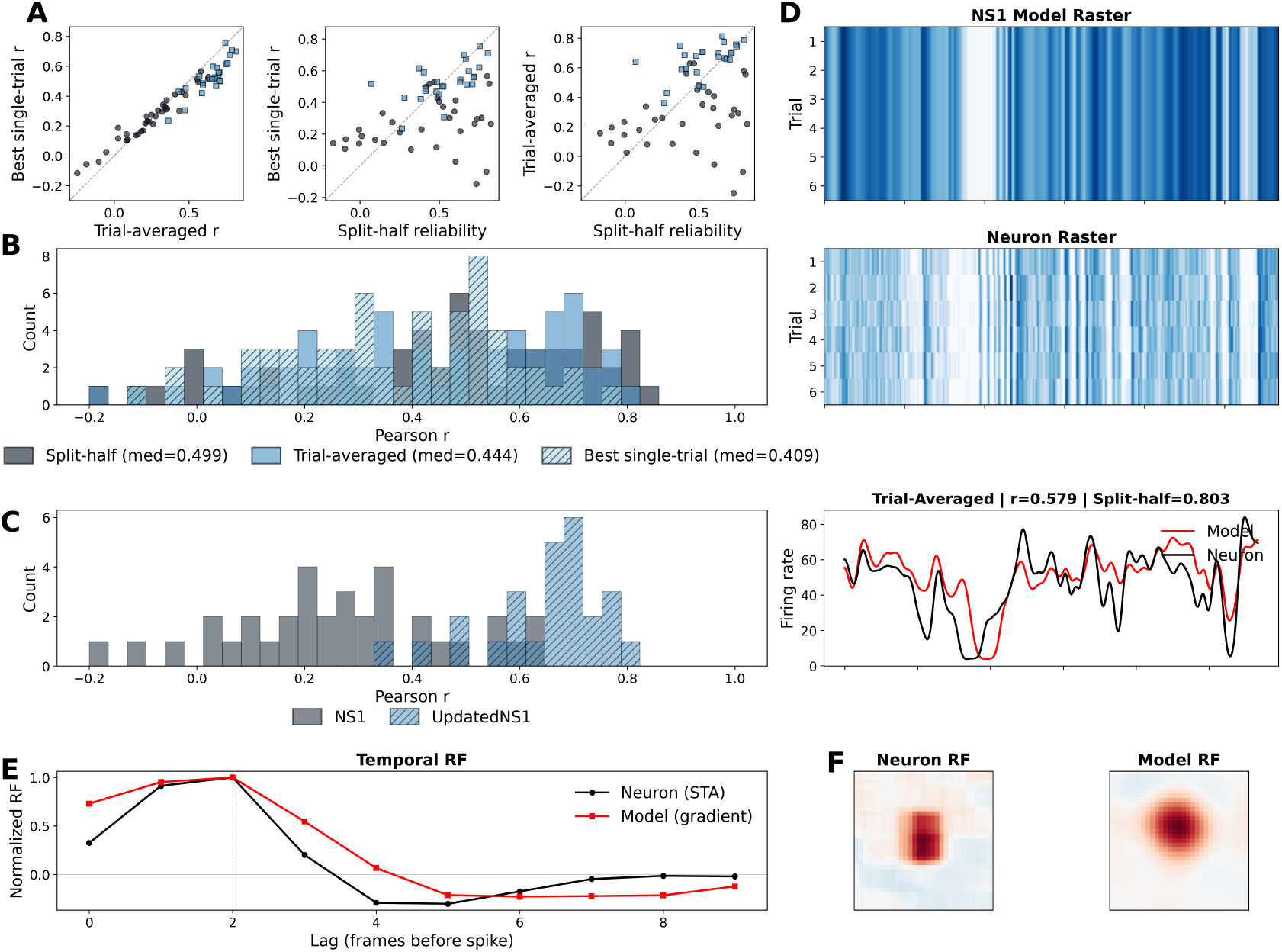
Model evaluation metrics for retinal extracellular single-cell recordings. **(A)** Single-trial, trial-averaged, and split-half reliability performance for predicting neural responses across two CNN models; different symbols indicate different animals. **(B)** Example model–neuron evaluation on a single trial and on the trial-averaged prediction (time scale in seconds). **(C)** Example neuron spike-triggered average (left) and model gradient spatial receptive field (right), measured with white noise.

**Fig. S3.**
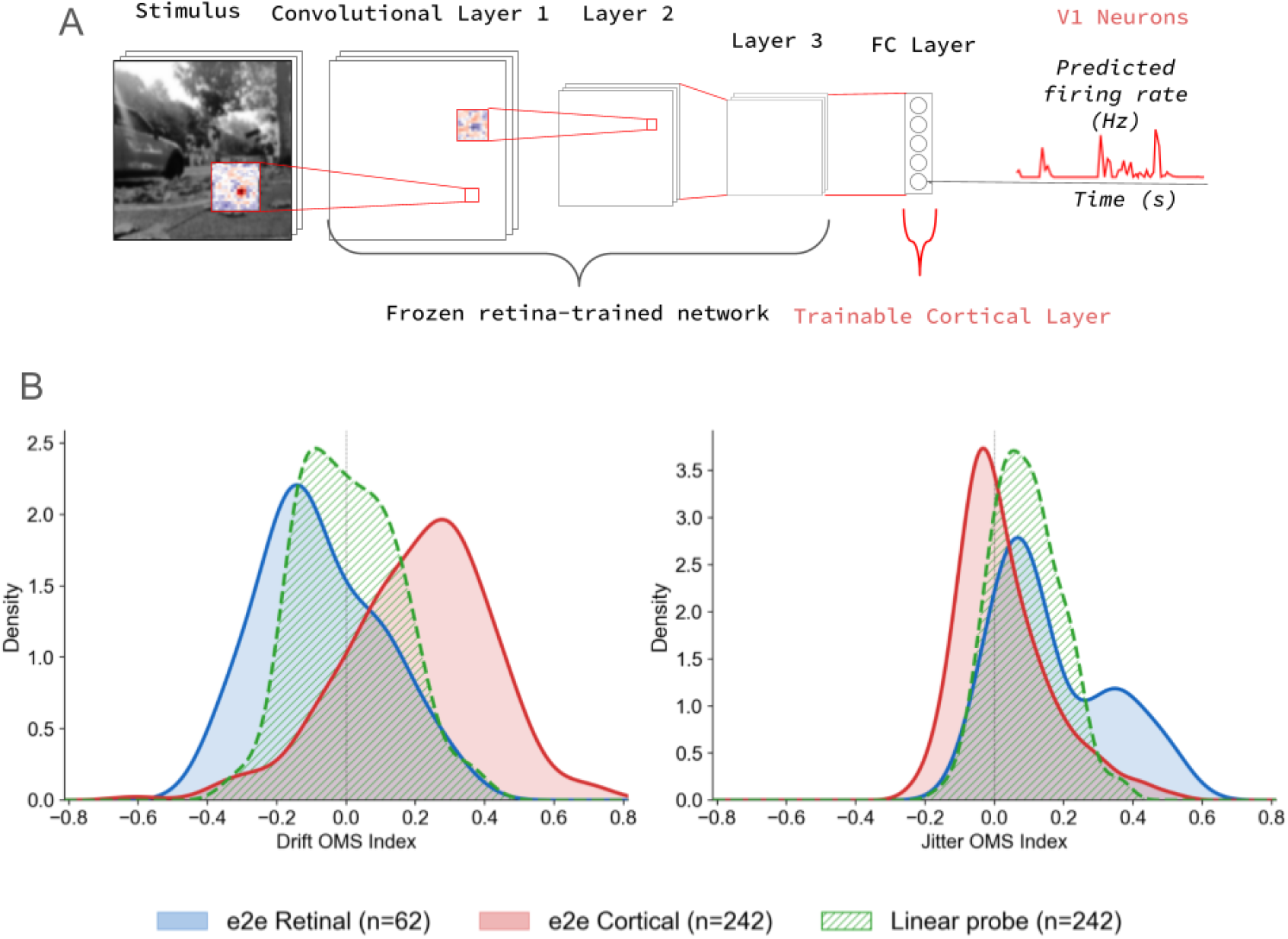
Trained retinal-cortical linear probe testing phenomenological recovery of the drift-selective OMS property found in the cortex. **(A)** Schematic of training a retinal linear probe with a frozen retina from end and linear readout layer to assess the linear transformation between the retina and cortex. **(B)** Drift vs. jitter OMS index (OMSI) for retinal and cortical models trained end-to-end (left), tested with identical OMS stimuli. Retinal units (blue) are jitter-preferring; cortical units (red) are drift-preferring. Stars, population means.

**Fig. S4.**
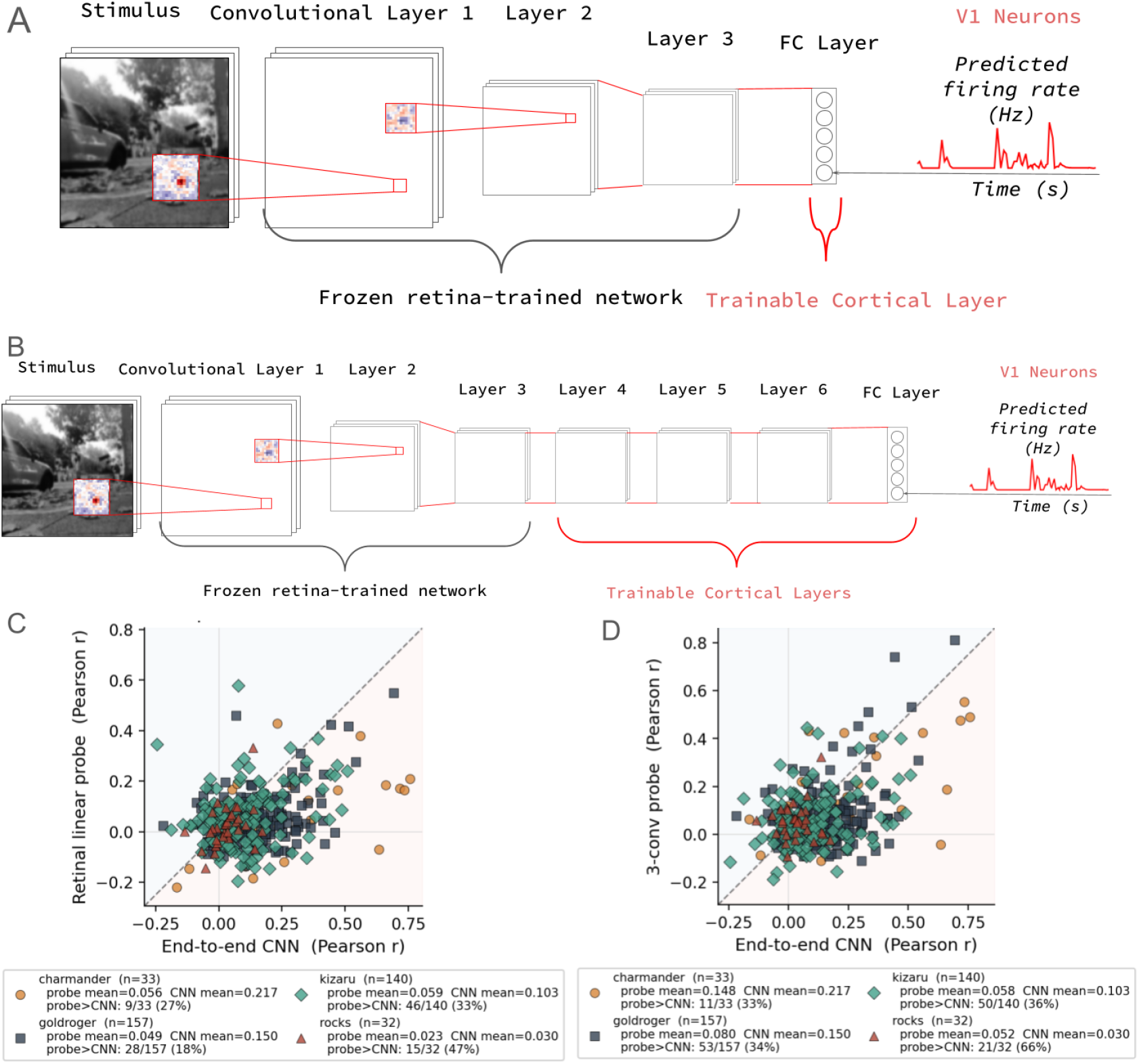
Single-trial firing rate prediction for linear probe (left) and 3Conv probe showing most cells are better predicted by the model trained end-to-end on V1 cortical data than constrained with retinal representations at the front end. **(A)** Schematic for linear probe of frozen trained retina front-end and a single linear layer testing linear transformability of V1 responses from retinal outputs. **(B)** Schematic for 3-layer-convolutional probe of frozen trained retina front-end and three randomly initialized convolutional layers with three additional nonlinearities and a fully connected layer testing predictability of V1 responses using retinal outputs as the bottleneck for training. **(C)** Results from the linear probe compared to the end-to-end model trained on the cortex show that most units are better predicted on a held-out, single-trial natural scenes repeat by a model trained on V1 end-to-end rather than through constraining a retinal front-end. **(D)** Results from the 3Conv probe compared to the end-to-end model trained on the cortex show that most units are better predicted on a held-out, single-trial natural scenes repeat by a model trained on V1 end-to-end rather than using the retina as the frozen front end. This probe shows marked improvement over the linear probe showing that multiple stages of nonlinearity are needed to recover some of the variance observed in V1 responses to natural scenes.

**Fig. S5.**
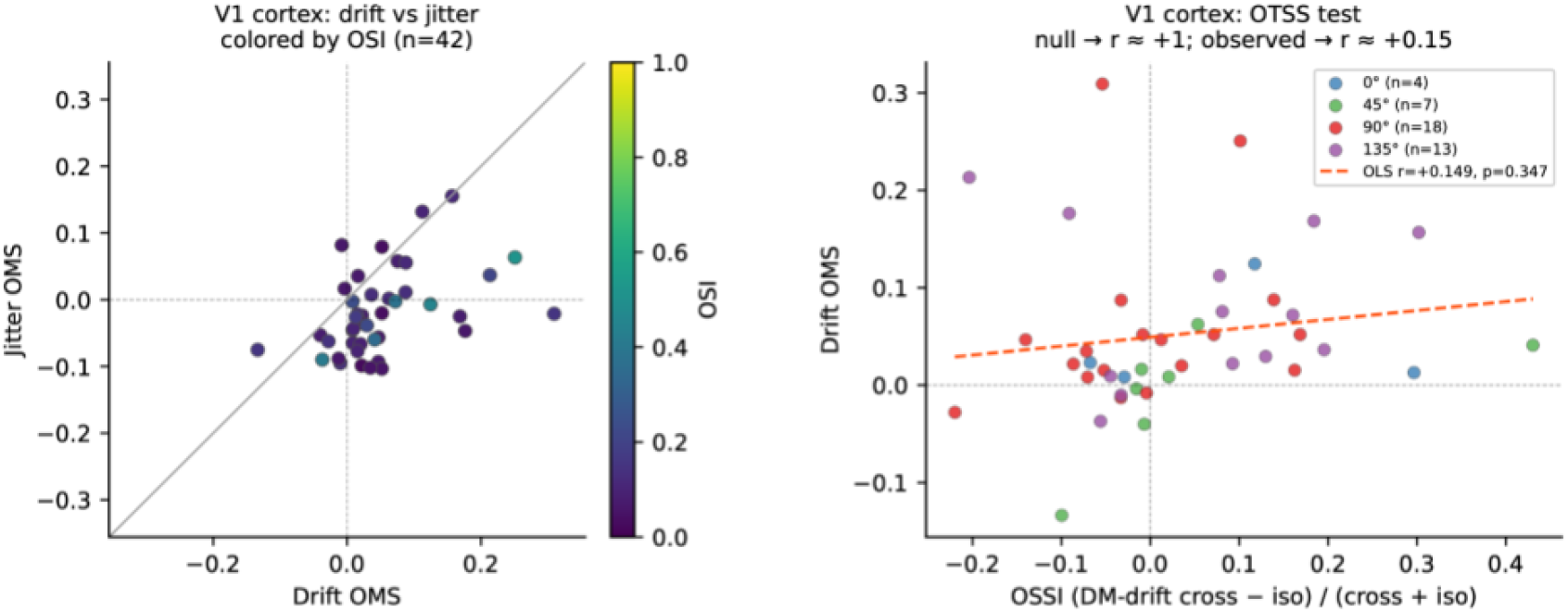
Test for orientation tuned surround suppression in cells identified as OMS cells by the OMSI in the targeted V1 experiment, showing variable orientation selectivity in recorded neurons and weak correlation with oriented surround suppression suggesting OTSS is weakly correlated with OMS.

**Fig. S6.**
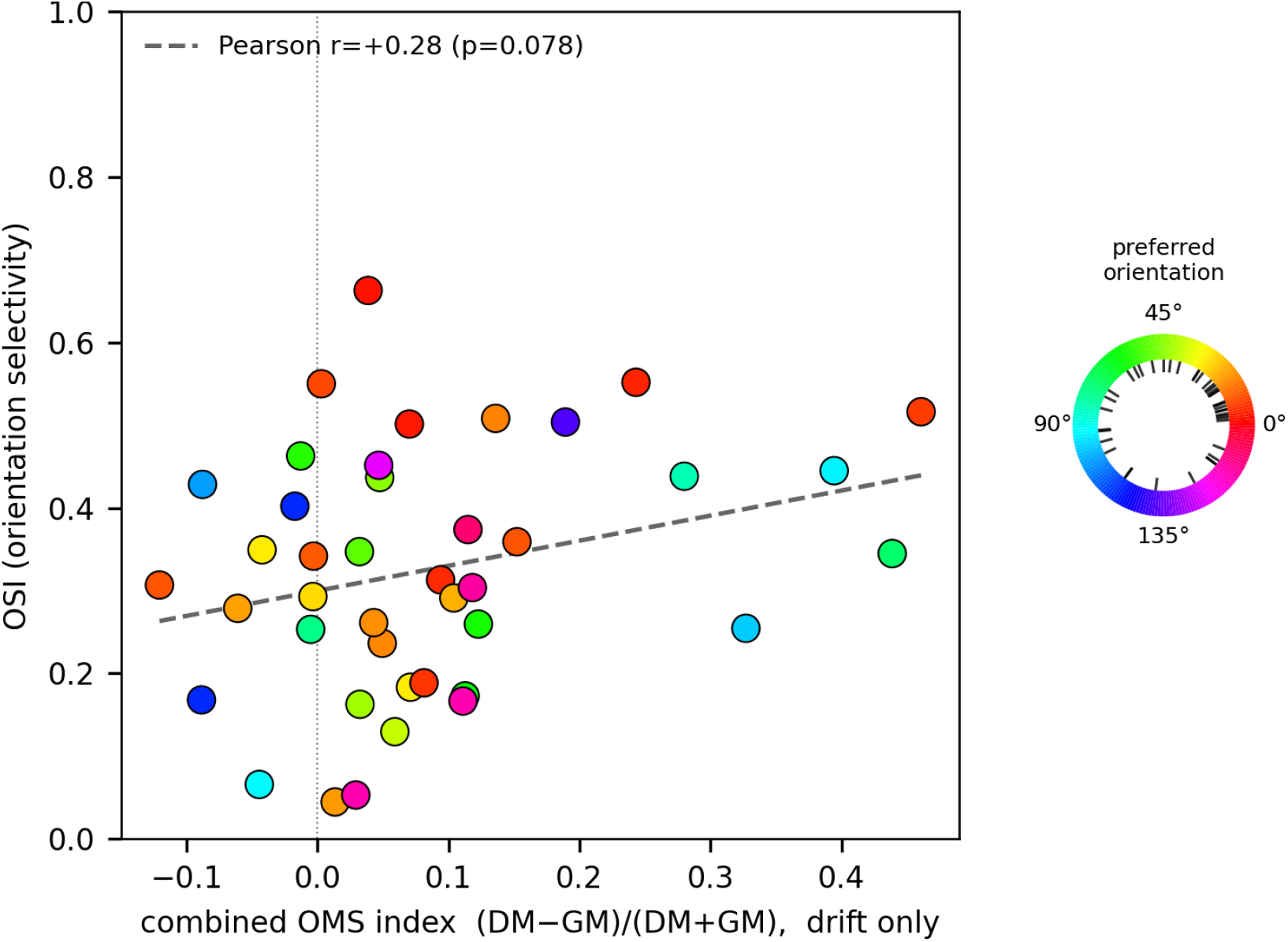
V1 OMS cells from targeted in-vivo recording: OMS strength vs OSI (n=41). There is a weak and statistically insignificant correlation between orientation selectivity and object motion sensitivity, showing that OMS is a computation that is orientation gated, but not fully driven by the orientation selectivity of the neurons in V1.

**Fig. S7.**
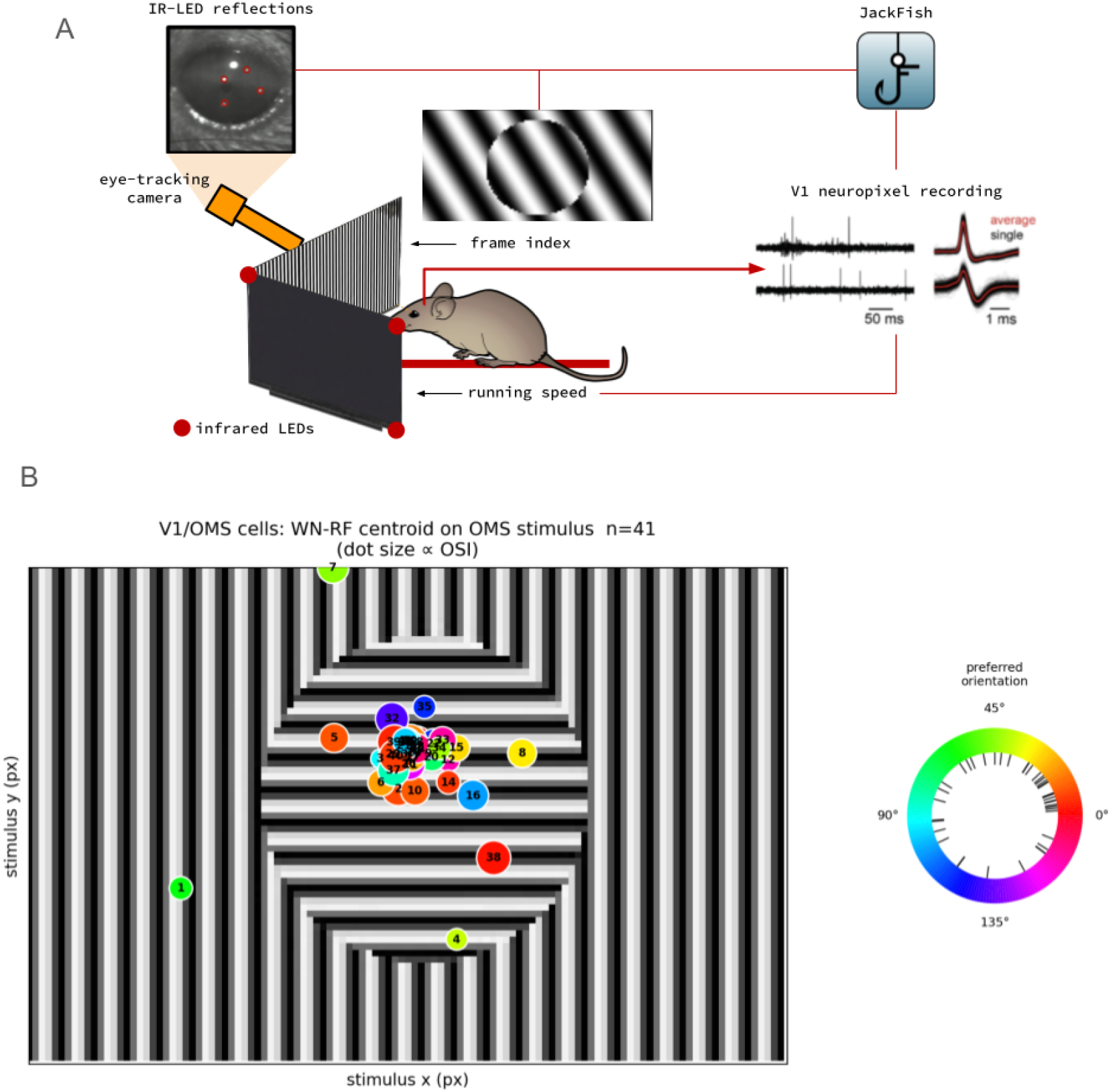
Receptive Field centroid and the orientation preferences of recorded V1 neurons during the targeted OMS experiment. Cells in the surrounding vicinity of the differential motion stimulus are not considered as they do not meet the centered criterion. **(A)** Schematic of targeted experiment testing whether OMS predicted in the model was present in the mouse primary visual cortex (V1). **(B)** The position of V1 OMS cells linear receptive field as measured by white noise spike-triggered averages. Colors indicate the orientation selectivity of the numbered cell.

**Fig. S8.**
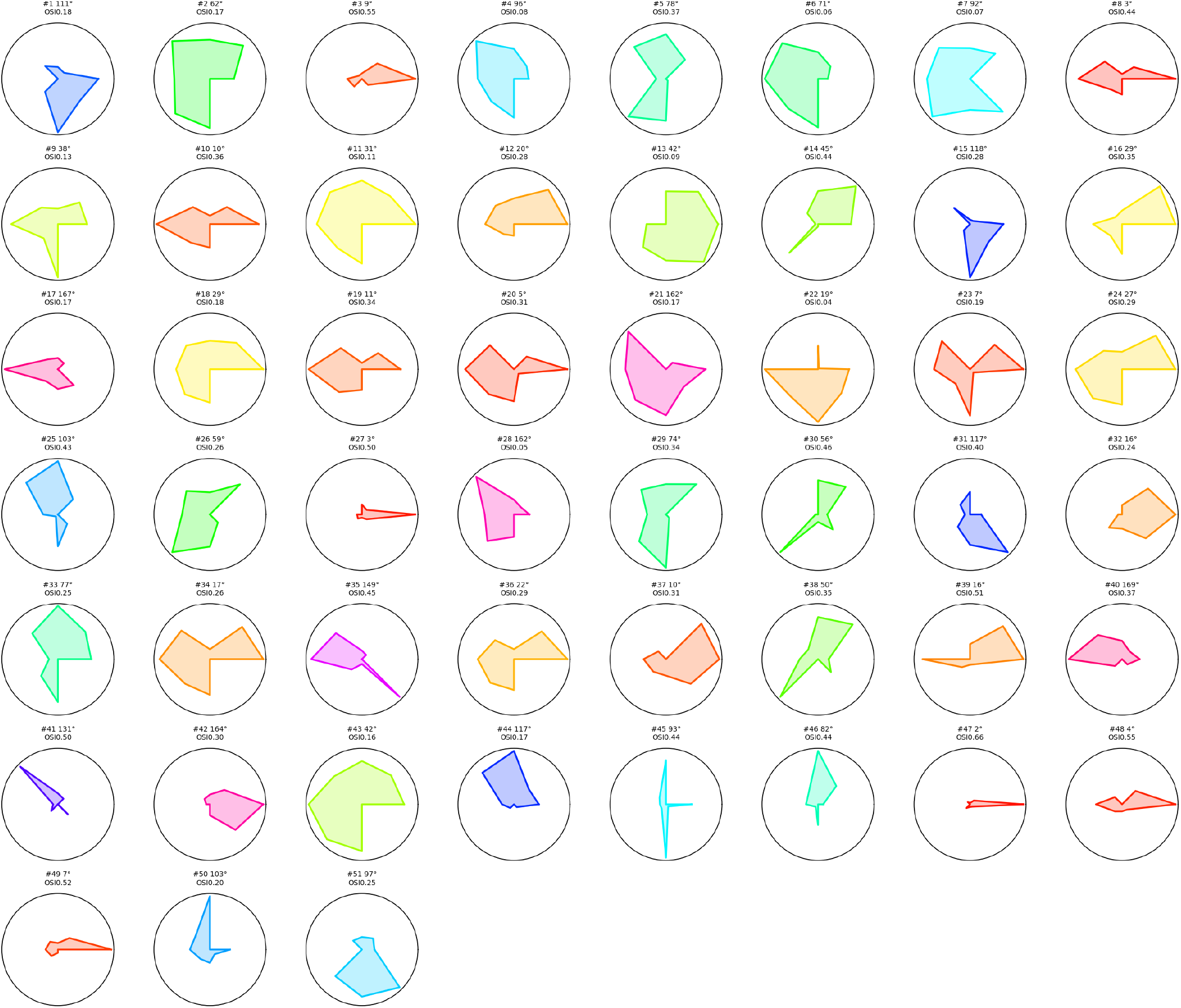
Full-field grating orientation tuning of the full population for the targeted V1 OMS experiment.

**Fig. S9.**
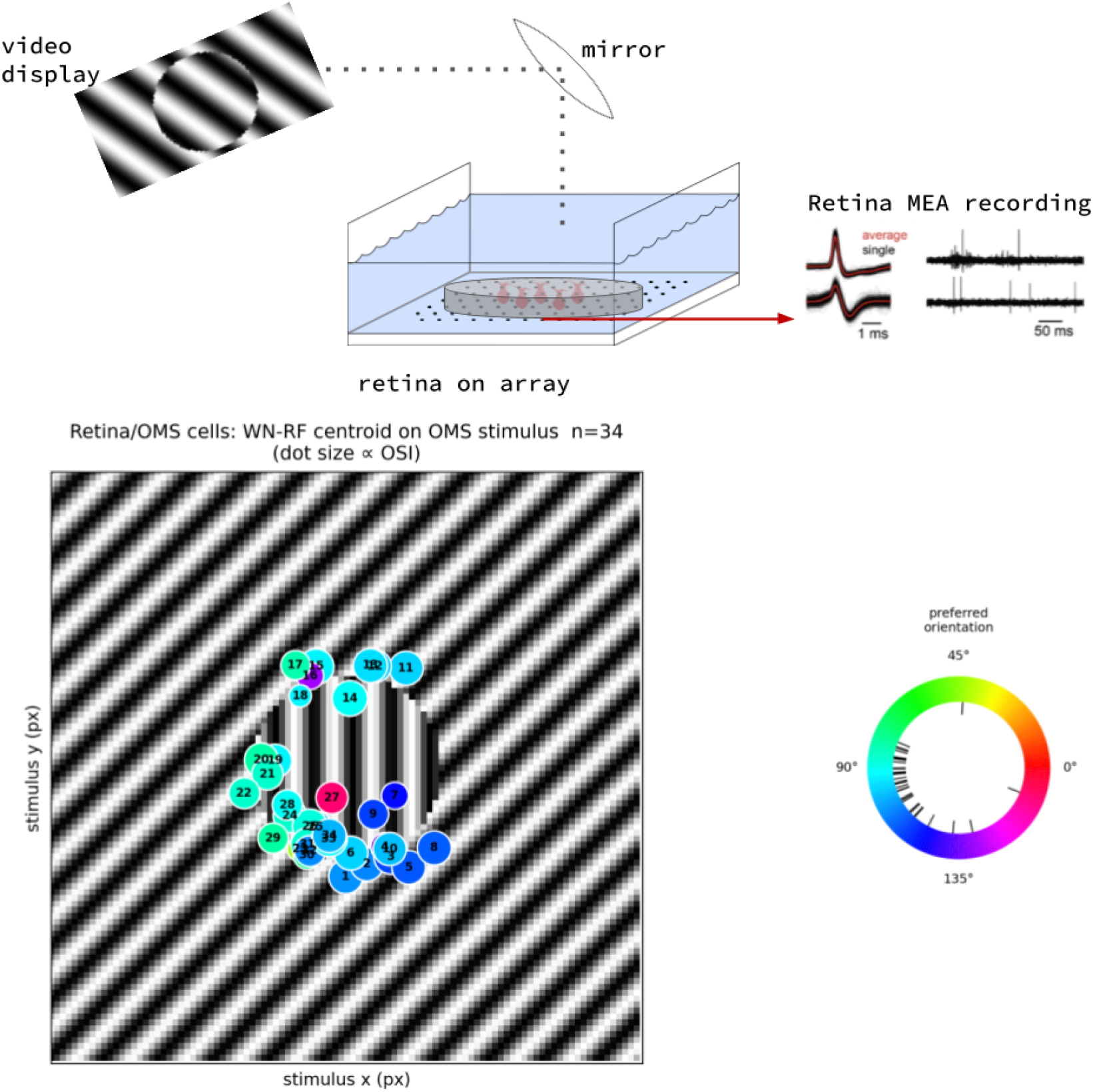
Receptive Field centroid and the orientation preferences of recorded ex-vivo retinal ganglion cell neurons during the targeted OMS experiment. Cells in the surrounding vicinity of the differential motion stimulus are not considered as they do not meet the centered criterion. **(A)** Schematic of targeted experiment confirming prior OMS in the retina. **(B)** The position of the retinal ganglion cell linear receptive field as measured by white noise spike trigger averages. Colors indicate the orientation selectivity of the numbered cell.

**Fig. S10.**
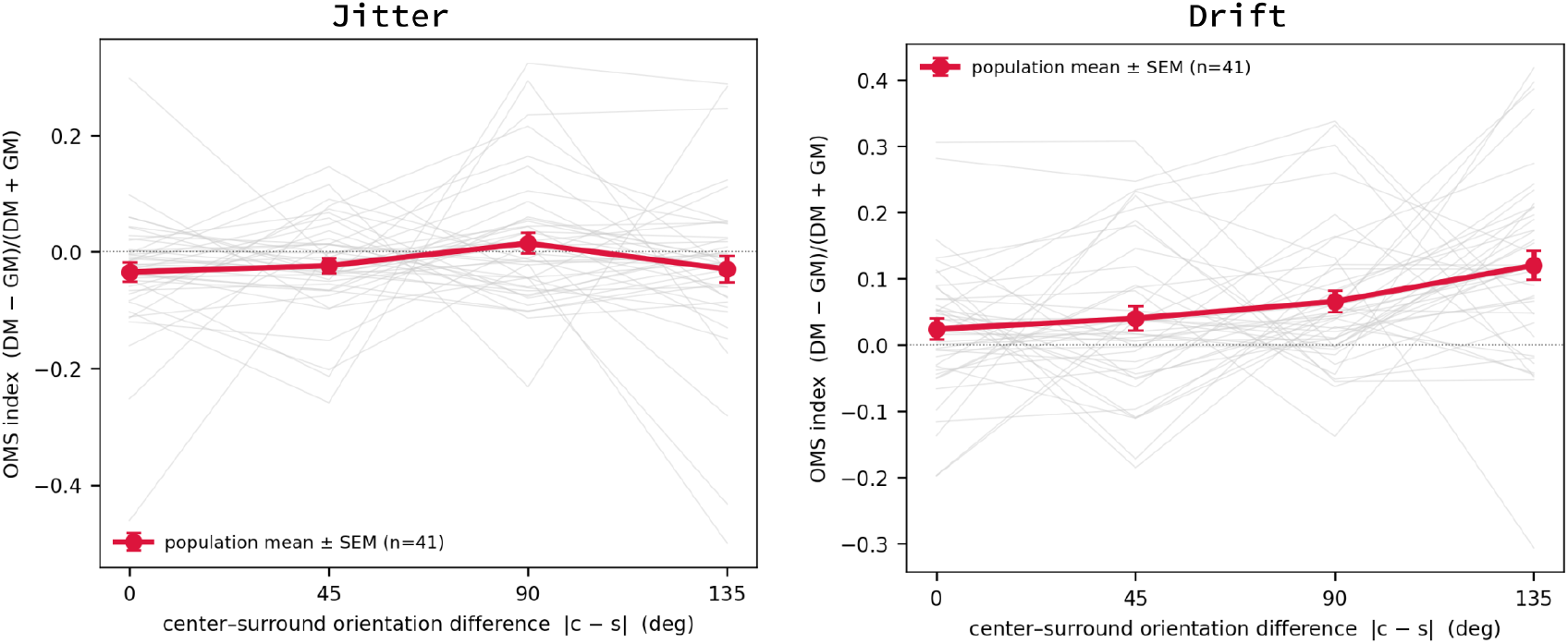
Orientation-gating of the OMS response in recorded V1 neurons depends on the relative orientation of the center and surround gratings. The OMS index was computed separately at four bins of center-surround orientation differences (0, 45, 90, 135). The four-point orientation tuning curve is plotted for every individual cell (grey lines) and their population mean (red lines). The orientation gated OMS response in the right panel showed a monotonic increase in for drifting differential per 90 degrees for drifting differential motion (*p <* 0.001), unlike the jitter OMS response where there is no significant orientation dependence (*p* = 0.36).

**Fig. S11.**
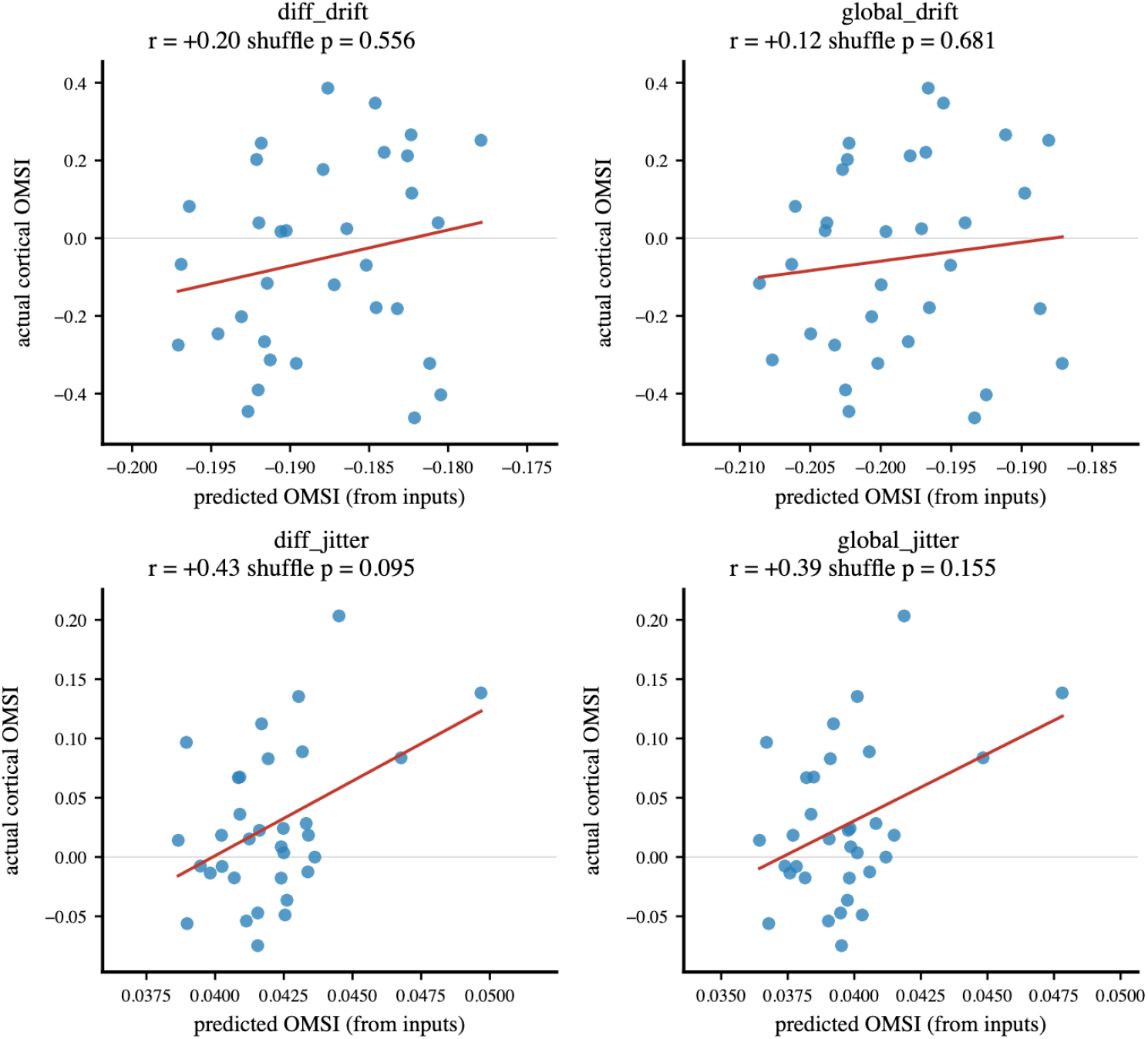
Object Motion Selectivity Index (OMSI) evaluated via a 3-convolutional-layer probe. We evaluate whether cortical Object Motion Selectivity (OMS) is inherited directly from retinal inputs across four motion conditions (differential drift, global drift, differential jitter, and global jitter) for n = 33 cortical cells. The analysis evaluates whether the cortical response is recoverable using a probe model where a fully convolutional network trained on the retina is frozen, followed by three randomly initialized trainable convolutional layers and a fully connected layer trained to predict V1 responses to natural scenes. The retinal OMSI is calculated as a weighted average of the OMSI from 26 retinal channels. The weights correspond to each channel’s absolute (excitatory plus inhibitory) contribution to the cortical cell, which was quantified using the Integrated Gradients method. Overall, the retinal channels’ OMSI does not strongly predict the actual cortical OMSI, supporting the hypothesis that the cortex primarily computes its own motion preferences rather than passively inheriting them. Specifically, drift stimuli demonstrate a null result for inheritance, yielding no significant correlation between predicted and actual values for either differential (r = +0.20, shuffle p = 0.556) or global drift (r = +0.12, shuffle p = 0.681). Jitter conditions exhibit slightly higher, though statistically weak, correlations (differential: r = +0.43, p = 0.095; global: r = +0.39, p = 0.155), suggesting a potential, minor inherited component for jitter preferences that is absent for drift stimuli.

**Table S1.** Correlation between predicted and actual cortical OMSI partitioned by excitatory and inhibitory retinal contributions. Object Motion Selectivity (OMS) inheritance by separating the retinal channel contributions into Excitatory (E), Inhibitory (I), and combined (E+I) components. For each of the four motion conditions (differential drift, global drift, differential jitter, and global jitter), the predicted cortical OMSI was calculated as a weighted average of the 26 retinal channels’ OMSI. The weights were determined by the magnitude of the specific contribution type (E, I, or E+I) derived via the Integrated Gradients method.

| Condition | Type | Correlation ($r$) | Shuffle $p$-value |
| --- | --- | --- | --- |
| diff_drift | E | +0.122 | 0.7004 |
| diff_drift | I | +0.055 | 0.8590 |
| diff_drift | E+I | +0.198 | 0.5560 |
| global_drift | E | +0.035 | 0.8850 |
| global_drift | I | +0.090 | 0.7004 |
| global_drift | E+I | +0.115 | 0.6812 |
| diff_jitter | E | +0.105 | 0.8102 |
| diff_jitter | I | +0.453 | 0.0548 |
| diff_jitter | E+I | +0.432 | 0.0946 |
| global_jitter | E | +0.063 | 0.8762 |
| global_jitter | I | +0.439 | 0.0528 |
| global_jitter | E+I | +0.394 | 0.1554 |

**Supplementary Videos (dynamic figures)**. These dynamic figures cannot be embedded in a static PDF. They are provided as *ancillary files* with the arXiv source package (in the `anc/` directory) and are listed for download on the arXiv abstract page. They can also be viewed in the interactive platform accompanying this paper at https://convergent-oms.github.io.

**Supplementary Video 1** (ancillary file `VideoS1.gif`). Representative frames from the MoCA dataset showing an arctic fox (left) with retinal (center) and cortical (right) heatmaps; the black bounding box identifies the ground-truth location. While retinal units respond with focal, motion-invariant signals, cortical population activity exhibits an extended, orientation-selective architecture that spans the body of the moving predator. We simulate fixational eye-movements by adding a sub-pixel random-walk jitter to the input (Gaussian steps of *σ* = 0.15 px ≈ 0.15·d° per frame, bounded to ≤ ±1 px ≈ ±1·d°).

**Supplementary Video 2** (ancillary file `VideoS2.gif`). Orientation and Direction quiver plots for the population vector decoding of orientation and direction of OMS and Non-OMS cells. OMS units are individually biased toward direction over orientation (DSI *>* OSI, *p* = 0.002 paired), whereas non-OMS units are not (*p* = 0.76); the two populations do not differ in orientation selectivity (*p* = 0.43) but OMS units are more direction-selective (*p* = 0.003).

**Fig. S12.**
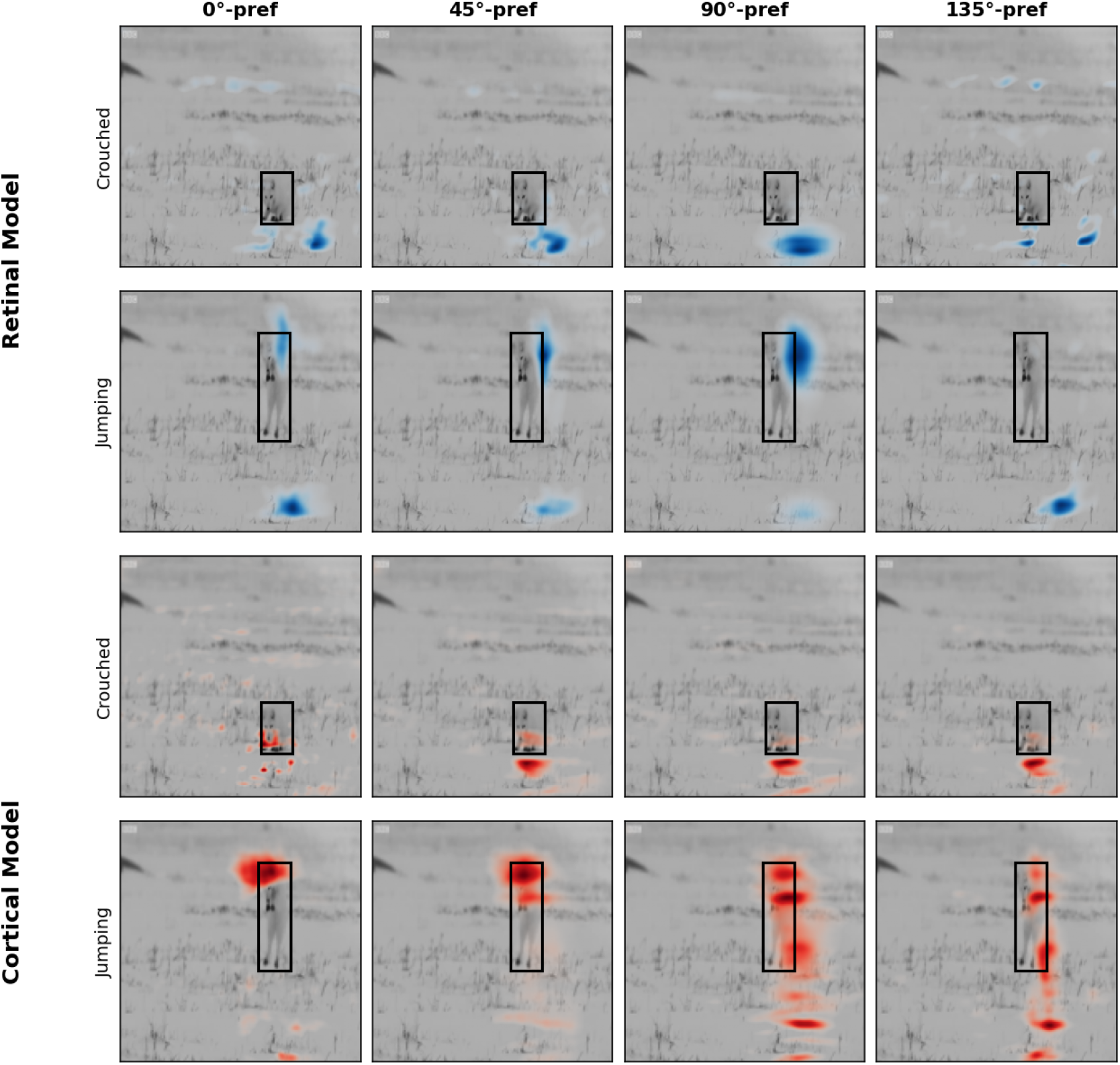
Channel-wise orientation preference read out of camouflaged motion detection in the retinal and cortical models, during stillness and locomotion. Activation of the arctic fox is shown for each of the units in the model; the same as in figure 6, grouped into four channels by each units’ preferred orientation (columns: 0, 45, 90, 135; measured with drifting object motion gratings). Rows show the two models (Retinal Model, top blue heatmap activations; Cortical Model, bottom red heatmap activations), both at two moments of the sequence: crouching and jumping. Activation is overlaid on the video frame with the black box marking the annotated region of the arctic fox. During object (fox) locomotion, both model’s oriented channels activate following the moving fox, confirming that each detects that an object is in motion. The models differ in how they represent object motion. The retina shows a compact focus that marks the animal’s presence but is largely invariant to the object’s extent, whereas the cortical channels tile along the fox’s elongated silhouettes; with the vertical (90 and 135 degree) channels tracking the length of the reared body. Thus the retinal model signals spatially invariant object motion, while the cortical model adds spatial granularity, resolving where along the animal’s body the motion occurs.

