## Supplementary figures and images for "Parallel construction of object motion from retina to cortex"

### Supplementary Video 1

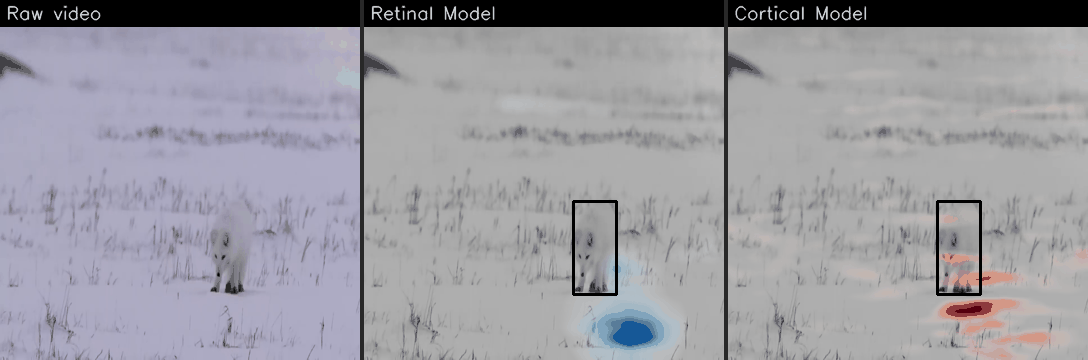

### Supplementary Video 2

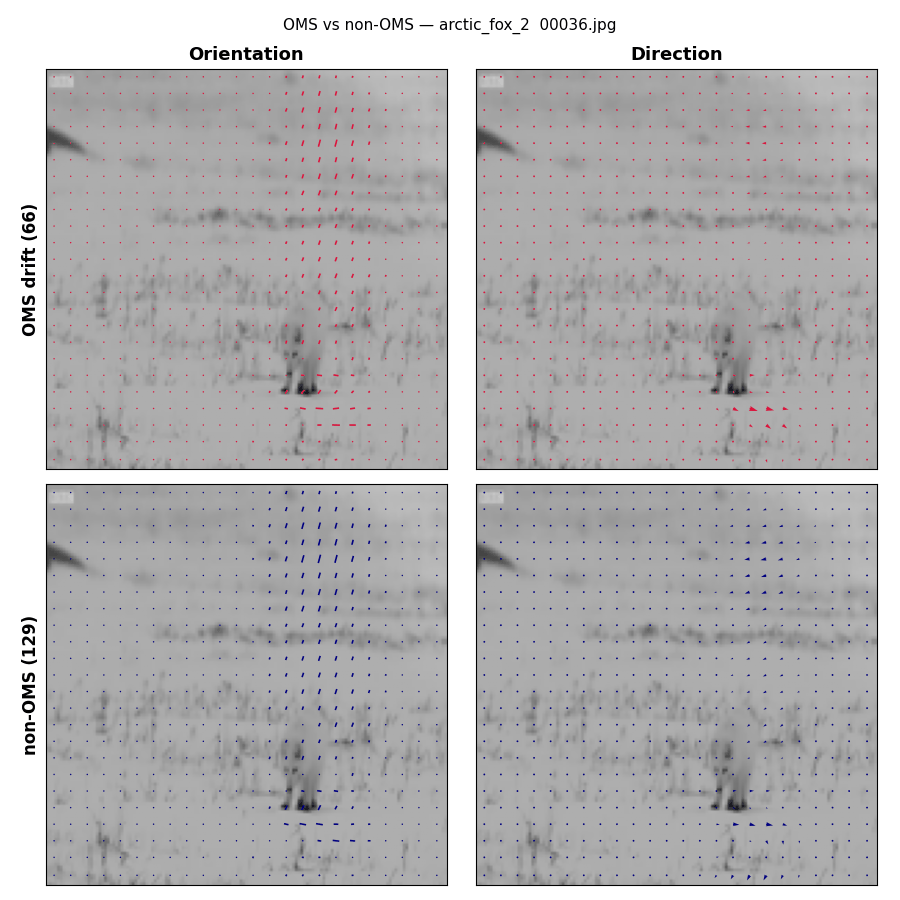
